# Chloroplast expression of *Chlamydomonas* glycolate dehydrogenase en route to an improved photorespiratory bypass

**DOI:** 10.64898/2026.07.23.739952

**Authors:** Jooyeon Jeong, Kwangryul Baek, Sherinmol Thomas, Samantha S. Stutz, Jeffrey M. Staub, Doug K. Allen, Yong-Su Jin, Donald R. Ort

## Abstract

Successes with photorespiratory bypass pathway engineering have relied on nuclear transformation, requiring subcellular targeting and protein localization in the target organelles for photorespiration. In the current study, mitochondrial *Chlamydomonas* glycolate dehydrogenase (CrGDH) was directly expressed in chloroplasts of *Chlamydomonas* and tobacco, as the first enzymatic step for a photorespiratory bypass. Proof-of-concept experiments in *Chlamydomonas* were followed by transgenic tobacco lines that confirmed the chloroplast accumulation of active CrGDH protein. Photosynthetic rates and biomass were comparable or less than those of wild type plants under the tested conditions, indicating that chloroplast expression of CrGDH alone is insufficient to improve plant performance. These results suggest that additional downstream enzymes within a complete photorespiratory bypass are required to metabolize glyoxylate derived from CrGDH activity and thereby benefit growth. One of the chloroplast transformed lines (APP2882) accumulated CrGDH while maintaining wild type levels of photosynthesis and biomass, providing a best candidate chassis for engineering full photorespiratory bypass pathways.

## Introduction

With increasing global population and the challenges to agriculture posed by climate change, achieving stable and sustainable crop yields has become an urgent priority (Long, 2025). Improving photosynthesis represents a promising strategy to enhance crop productivity, and within this context, improving photorespiratory efficiency has emerged as an attractive target in C3 plants (Bernacchi et al., 2025; Croce et al., 2024; Long et al., 2025). Photorespiration is a metabolic process initiated by the oxygenation of ribulose-1,5-bisphosphate by Rubisco in the chloroplast, producing the two-carbon compound phosphoglycolate that is inhibitory to certain photosynthetic enzymes (Anderson, 1971; Flügel et al., 2017; Kelly & Latzko, 1976) necessitating its removal from chloroplasts. Multiple cellular compartments, including cytosol, chloroplasts, peroxisomes, and mitochondria, are involved in this energetically expensive process to salvage a portion of the photosynthetically fixed carbon and direct it back into the photosynthetic cycle. In C3 plants, the energy loss attributed to photorespiration has been estimated to account for approximately 6.1% of incident solar energy that could otherwise be utilized for photosynthesis (Zhu et al., 2010). Considering that the theoretical maximum efficiency of converting solar radiation into biomass in C3 plants is 4.6%, this loss represents one of the major limitations on photosynthesis and highlights photorespiration as a promising target for yield improvement (Walker et al., 2016; Zhu et al., 2010). Indeed, the introduction of synthetic pathways bypassing native photorespiration has often resulted in biomass and yield increases in various plant species (G. Chen et al., 2025; Dalal et al., 2015; Kebeish et al., 2007; Li et al., 2024; Lin et al., 2025; Maier et al., 2012; Roell et al., 2021; Shen et al., 2019; Wang et al., 2020; Xu et al., 2023). In our previous work, expression of photorespiratory alternative pathway 3 (AP3) in tobacco and potato resulted in enhanced yield gains, particularly under heat stress conditions that exacerbate photorespiration (Cavanagh et al., 2022; Meacham-Hensold et al., 2024; South et al., 2019).

Most of the photorespiratory bypass pathways implemented to date begin with the conversion of glycolate to glyoxylate in the chloroplast (G. Chen et al., 2025; Kebeish et al., 2007; Maier et al., 2012; Shen et al., 2019; South et al., 2019; Wang et al., 2020; Xu et al., 2023). Transgenes to catalyze this conversion have been introduced into the nuclear genome, which are then translated in the cytosol and subsequently imported into the chloroplast. Thus, each transgene must be equipped with an efficient chloroplast transit peptide (CTP) appropriate for the host species (Eseverri et al., 2020) and the cargo protein (Caspari, 2022). However, even if a CTP is derived from the same species and is known to be functional, targeting efficiency can vary depending on the protein structure of the transgene and the length of the CTP, and will likely require multiple rounds of optimization (Shen et al., 2017).

Direct transformation of the transgene into the chloroplast genome offers several potential advantages. Chloroplast transformation does not require a CTP and stable transgene insertions face fewer risks of gene silencing or positional effects because plastid transformation relies on homologous recombination targeted to a specific site in the plastid genome (Bock, 2015). Moreover, the high copy number of the chloroplast genome results in greater protein accumulation (Hanson et al., 2013), which could avoid bottlenecks at specific metabolic steps (Xin et al., 2014). Chloroplasts are inherited maternally thereby avoiding transgene escape via pollen (Bock, 2015). Chloroplast transformation is a promising route toward overcoming the glycolate to glyoxylate conversion bottleneck that constrains photorespiratory bypass designs.

In this study, we employed chloroplast transformation to overexpress *Chlamydomonas reinhardtii* glycolate dehydrogenase (CrGDH), which was used in the AP3 bypass (South et al. (2019) to convert glycolate to glyoxylate. As an initial rapid proof of concept, we first introduced mitochondrial CrGDH into the chloroplast of *Chlamydomonas cia5* CRISPR-Cas9 mediated knockout mutant and observed a reduction in glycolate accumulation. We then introduced CrGDH into tobacco chloroplasts by chloroplast transformation, and confirmed its localization to the thylakoid membrane, and quantified its expression and enzymatic activity. This transgenic tobacco line is intended to serve as a transformation chassis for construction of AP3 and other bypasses that utilize CrGDH. Furthermore, because the overexpression of *Escherichia coli* glycolate dehydrogenase (EcGDH) has unexpectedly been reported to confer photosynthetic and growth advantages without presence of additional bypass genes (Dalal et al., 2015; Kebeish et al., 2007; Nölke et al., 2014), we evaluated the expression of CrGDH on photosynthesis and growth in tobacco.

## Materials and Methods

### Generation of Chlamydomonas transgenic strains

*Chlamydomonas reinhardtii* CC-4349 (*cw15 mt*−) was used as the parental strain for the generation of transgenic lines. CRISPR-Cas9-mediated knock-in targeting *CIA5* gene (Cre02.g096300) was performed based on the method described by Baek et al. (2016), with minor modifications. The *CIA5*-targeting sgRNA and Cas9 protein were prepared according to Yu et al. (2017). The hygromycin resistance cassette used for knock-in was amplified by PCR using GeneArt pChlamy3 plasmid (Invitrogen) as a template. The primers used are listed in Supplementary table S1. For knock-in transformation, a total of 5 × 10^5^ cells were used. Cells were mixed with the ribonucleoprotein complex, consisting of 100 µg Cas9 protein and 70 µg sgRNA synthesized by *in vitro* transcription, together with 1 µg knock-in DNA. The mixture was introduced into cells using Bio-Rad Gene Pulser Xcell Electroporation System following the recommended protocol of GeneArt Chlamydomonas Engineering Kit (Invitrogen). After transformation, cells were plated on TAP medium containing 1.5% agar and 50 µg ml^-1^ hygromycin to obtain single colonies. The resulting single colonies were initially screened by colony PCR targeting the *CIA5* locus, following the method described by Cao et al. (2009). The primers used for colony PCR are listed in Supplementary table S1. Disruption of the *CIA5* gene was further confirmed by Sanger sequencing. The confirmed *cia5* knockout lines were subsequently used for immunoblot analysis and glycolate quantification experiments.

Among the selected *cia5* knockout lines, the *cia5* KO 1 strain was used as the recipient strain for chloroplast transformation to generate CrGDH-overexpressing lines. Prior to the introduction of CrGDH, a *psbH* knockout line was generated to enable selection under photoautotrophic conditions, using a modified version of the method described by Economou et al. (2014) and Young and Purton (2016). Briefly, the *psbH* gene and empty expression cassette from the pSRSapI plasmid were replaced by spectinomycin resistance cassette from the pWUCA1 plasmid, and the resulting construct was introduced into chloroplasts and integrated into the single copy region between *Chlamydomonas trnE2* and *psbN* genes. This resulted in the deletion of the *psbH* gene and conferred spectinomycin resistance simultaneously. Chloroplast transformation was performed using the PDS-1000/He particle bombardment system (Bio-Rad). Plasmid DNA was coated onto 0.6 µm gold microcarriers (Bio-Rad) and delivered into cells by bombardment. For each bombardment, 7.5 µg plasmid DNA was used. DNA coating was performed using 2.5 M CaCl_2_ and 0.1 M spermidine. To facilitate the recovery of homoplasmic transformants, cells were treated with 0.5 mM 5-fluoro-2′-deoxyuridine for 5 days prior to transformation. A 10 µl cell suspension containing 2 × 10^6^ cells was plated onto TAP medium containing 1.5% agar and 100 µg ml^-1^ spectinomycin. Bombardment was performed using a 1,100 psi rupture disk, a vacuum pressure of 28 in Hg, and a target distance of 6 cm. After bombardment, cells were incubated at 25 °C under dim light conditions, approximately 5 µmol photons m^-2^ s^-1^. The resulting transformants were grown under both photoheterotrophic and photoautotrophic conditions. Colonies that grew under photoheterotrophic conditions but failed to grow under photoautotrophic conditions were selected as candidate *psbH* knockout lines. The disruption of both *CIA5* and *psbH* was confirmed by PCR and Sanger sequencing (Supplementary table S1).

The confirmed *cia5 psbH* knockout line was then used for a second round of chloroplast transformation with the pSRSapI-CrGDH plasmid, which contains both *CrGDH-HA* and an intact *psbH* gene. The *psbH* gene was cloned at its native position between *Chlamydomonas trnE2* and *psbN* genes using 1,946 nts and 800 nts flanking sequences. The *CrGDH* gene (Cre06.g288700) was codon-optimized for chloroplast expression and cloned 151 nts downstream of *<u>psbH</u>* in the reverse orientation. The *CrGDH* was fused to a C-terminal 3×HA epitope tag and was driven by *Chlamydomonas psbA* gene promoter (P*psbA*) and *rbcL* gene 3’-end (T*rbcL*). Transformants were selected under photoautotrophic conditions, and successful integration of *CrGDH* in the *cia5* knockout background was confirmed by PCR and Sanger sequencing.

Cells were grown at 25 °C under continuous light at 100 µmol photons m^-2^ s^-1^. Photoheterotrophic cultures were grown in Tris-acetate-phosphate (TAP) medium, whereas photoautotrophic cultures were grown in high-salt (HS) medium.

### Glycolate quantification

Glycolate released into the culture medium by *Chlamydomonas* strains grown under photoautotrophic conditions was quantified by high-performance liquid chromatography (HPLC). Culture samples were centrifuged at 20,000 × g for 5 min at room temperature, and the resulting supernatants were filtered through a 0.22 µm syringe filter. Filtered samples were analyzed using an Agilent Technologies 1200 Series HPLC system equipped with a Rezex ROA-Organic Acid H^+^ (8%) column (Phenomenex). The mobile phase was 0.005 N sulfuric acid, and the column temperature was maintained at 50 °C. The flow rate was 0.6 ml min^-1^, and the injection volume was 10 µl. Glycolate was detected using a refractive index detector, and its concentration was determined from a standard curve generated using glycolic acid standards (Sigma-Aldrich).

### Generation of tobacco transgenic lines

The CrGDH and GFP genes were synthesized using plastid-preferred codons according to previous methods (Staub et al., 2000). The CrGDH coding region differs from previously reported (South et al., 2019) by a E155D change, designed to remove a restriction enzyme site. For the CrGDH-GFP fusions, the GFP coding region (100% identity to amino acids 68 – 769 of GenBank OP654541.1) is separated from CrGDH by a GGVS amino acid linker. The HiBit tag (VSGWRLFKKIS) is fused directly to the C-terminus of CrGDH. The *aadA* gene used for selection of chloroplast transformants was driven by the tobacco P*rrn* promoter (Suzuki et al., 2003) and the *E. coli rrnB* terminator fragment (T*rrn*; J. Zhang et al. (2015)). The GFP and CrGDH coding regions were driven by the tobacco *psbA* gene promoter (P*psbA;* Staub and Maliga (1993)) and the *petD* gene 3’-end (T*petD*). In pAPP2881 and pAPP2882, the *aadA* and *CrGDH* transgenes are cloned in the Single Copy Region between and in the same transcriptional orientation as the tobacco *rbcL* and *accD* genes, and are flanked by 1295 and 1150 nts, respectively, of identity to the tobacco (*N. tabacum* cv Petit Havana) chloroplast genome. In pPTS78, pAPP2883 and pAPP2884, the transgenes are cloned between the tobacco chloroplast *trnV* and *3’rps12/7* genes in the Inverted Repeat Region and are flanked by 987 and 1005 nts, respectively, of identity.

Plant growth, transformation, and selection of transformants was essentially as described previously (Bélanger et al., 2023). Briefly, tobacco plants were grown aseptically from seedlings on MS agar medium for ∼4 weeks at 28 °C under a 16 h light/8 h dark photoperiod. Young leaves were harvested for particle bombardment, placed abaxial side up and bombarded using the BioRad PDS1000 He gun. Transplastomic events were selected by growth on 500 mg/L spectinomycin. Young leaf tissue from primary transformants was dissected and used for a second and subsequently third round of plant regeneration on selective medium to ensure homoplasmy of the plastid transformed lines. Plastid transformants were confirmed by PCR-sequencing of amplification products to confirm the entire transgenic insert sequence, including correct junctional sequences with nontransformed wild-type chloroplast genome. As expected for homologous recombination in plastids, all independent chloroplast transgenics lines derived from a single transformation vector were molecularly identical. Consequently, a single line was used in some cases for detailed biochemical analysis (see below).

Homoplasmic lines were transferred to soil to set seed. Homoplasmy and maternal inheritance was confirmed by self and reciprocal crosses of plastid transformed lines to wild-type plants. Uniform spectinomycin resistance of maternal-derived T1 seedlings is diagnostic for homoplasmy, while bleaching indicates wild-type spectinomycin sensitive plastid genomes (Supplementary figure S1).

For all subsequent experiments, homoplasmic T1 plants were grown in 1-gallon pots with a 20 cm top diameter, filled with BM6 All Purpose potting mix (Berger) supplemented with controlled-release fertilizer at ∼20 g per pot (Osmocote; 15-9-12). Plants were maintained in a controlled environment growth chamber at 25 °C under a 16h light/8h dark photoperiod with a light intensity of 300 µmol photons m^-2^ s^-1^ measured at the leaf level and 50% relative humidity. Except for GFP fluorescence measurements and confocal microscopy, all experiments were conducted using 6-week-old plants.

### GFP fluorescence measurements and confocal microscopy

Leaf discs 0.5 cm in diameter were sampled from the youngest fully expanded leaves of 3-week-old tobacco and placed in a black 96-well microplate containing 50 μL of DW. GFP and chlorophyll fluorescence signals were measured using a SpectraMax Plate Reader (Molecular Devices) with excitation/cutoff/emission settings of 392/475/507 nm for GFP and 435/630/640 nm for chlorophyll. For imaging, fresh leaf tissues were mounted in perfluorodecalin on glass slides and observed using a Zeiss LSM 880 confocal microscope (Zeiss) equipped with a 40 × water-immersion objective (NA 1.2). GFP fluorescence was excited with a 488 nm laser and detected at 487-543 nm, and chlorophyll autofluorescence was excited with a 561 nm laser and detected at 645-701 nm. Images were acquired in line-sequential mode and analyzed using Zeiss Zen version 3.9 software (Zeiss). A total of 10-50 imaging fields from 3-5 independent leaf samples were collected per line.

### Extraction of total soluble protein, chloroplasts, and thylakoid membranes

For total soluble protein extraction, leaf discs 1.25 cm in diameter were sampled from the youngest fully expanded leaves of 6-week-old tobacco plants. The leaf discs were frozen in liquid nitrogen, disrupted using a TissueLyser (Qiagen) for 1 min at 20 s^-1^, and then mixed with 0.2 mL of extraction buffer (125 mM Tris-Cl, pH 8.0, 1% (w/v) SDS, 10% glycerol, 50 mM sodium metabisulfite, and protease inhibitor cocktail). The samples were incubated for 10 min at room temperature, centrifuged for 5 min at 20,000 × g at room temperature, and the supernatants were used as total soluble protein extracts. For intact chloroplast extraction, a chloroplast isolation kit (Sigma-Aldrich) was used according to the manufacturer’s protocol. Thylakoid membranes were extracted from intact chloroplasts as described in Bouchnak et al. (2018). Proteins were quantified using DC Protein Assay kit (Bio-Rad) according to the manufacturer’s protocol.

### Total protein staining and immunoblot analysis

Proteins were separated on 4-20% Mini-PROTEAN TGX precast gels (Bio-Rad) and transferred to 0.2 μm nitrocellulose membrane using Trans-Blot Turbo transfer system (Bio-Rad) according to the manufacturer’s protocol. Total proteins on membrane were stained using Revert 700 Total Protein Stain kit (LI-COR Biosciences, Lincoln, NE, USA) and detected using Odyssey CLx imaging system (LI-COR) according to the manufacturer’s protocol. For Immunoblot analysis, membranes were then probed with polyclonal antibodies against GFP (Invitrogen, catalog no. CAB4211, 1:1,000 dilution), HA tag (Cell Signaling Technology, catalog no. 3724, 1:1,000 dilution), and photosynthetic proteins (Anti-ATPA antibody: PhytoAB, catalog no. PHY3003S, 1:1,000 dilution; Anti-PsaB antibody: PhytoAB, catalog no. PHY2559A, 1:1,000 dilution; Anti-PetB antibody: PhytoAB, catalog no. PHY0020S, 1:1,000 dilution; Anti-cFBPase antibody: Agrisera, catalog no. AS04 043, 1:1,000 dilution; Anti-RbcL antibody: Agrisera, catalog no. AS03 037, 1:10,000 dilution; Anti-PsbA antibody: Agrisera, catalog no. AS05 084, 1:10,000 dilution) followed by Alexa Fluor Plus 800 conjugated secondary antibody (Invitrogen, catalog no. A32735, 1:10,000 dilution). Signals were visualized using Odyssey CLx imaging system (LI-COR). For HiBiT blot analysis, membranes were analyzed using Nano-Glo HiBiT blotting system (Promega) and detected using ImageQuant 800 (Cytiva) according to the manufacturer’s protocol. All images were analyzed using ImageJ 1.54i for deconvolution and quantification of protein bands.

### Absolute protein quantification via LC-MS

Total proteins from tobacco leaves were extracted from 6-weekd-old tobacco plants. using a modified phenol-based method described by (Isaacson et al., 2006). Briefly, frozen leaf tissue powder was homogenized in an extraction buffer containing 0.7 M sucrose, 0.1 M KCl, 0.5 M Tris-HCl (pH 7.5), 50 mM EDTA, 50 mM dithiothreitol (DTT), 1 mM phenylmethylsulfonyl fluoride (PMSF), and 25 µL protease inhibitor cocktail. An equal volume of Tris-saturated phenol was then added, and the mixture was shaken at 4 °C for 30 min. Following centrifugation at 13,000 × g for 30 min at 4 °C, the phenolic phase was collected, and proteins were precipitated at -80 °C by adding five volumes of 0.1 M ammonium acetate containing 50 mM DTT. The protein pellet was recovered by centrifugation (13,000 × g, 30 min, 4 °C), washed twice with methanol containing 10 mM DTT, and finally washed with acetone containing 10 mM DTT. The resulting pellet was dissolved in 8 M urea buffer for further analysis.

Protein concentration was determined using a BCA protein assay kit (Pierce) according to the manufacturer’s instructions. To generate the isolation list for the proteomics study in Skyline, the CrGDH sequence corresponding to Cre06.g288700 was used. The transitions used for method development and quantification are summarized in Supplementary table S2. The APP2881-7a line was used as a positive control to evaluate peptide performance, and the peptide GLIPIVGAAR was selected as the surrogate peptide for Cr-GDH quantification based on its performance, uniqueness, and stability in our preliminary analysis of the APP2881-7a protein sample (Supplementary table S3, S4 and Supplementary figure S2).

For sample preparation, 50 μg of protein from each sample was digested into peptides using trypsin-Lys C (1:20 w/w; Pierce) for 16 h at 37 °C, followed by vacuum drying. Subsequently, 15 fmol of AQUA QuantPro heavy peptide GLIPIVGAAR (Pierce) was spiked into each sample. The peptides were then cleaned using C18 desalting tips (Pierce), and the resulting peptides were dissolved in 0.1% formic acid for LC-MS analysis.

For LC-MS analysis, 1 μL of sample was loaded onto a reverse-phase C18 column (Acclaim™ PepMap™ 100 C18 HPLC Columns, 2 μm particle size) and separated using a nano-flow Dionex RSLCnano HPLC system coupled to an Orbitrap Fusion Lumos mass spectrometer (Thermo Fisher Scientific), operated in positive parallel reaction monitoring (PRM) mode.

All raw data files were imported and analyzed using Skyline Daily (MacCoss Lab, University of Washington) (Pino et al., 2020). Absolute abundance was determined based on the normalized peak areas of the spiked heavy peptide (Sohn et al., 2018). The mass spectrometry proteomics data have been deposited to the ProteomeXchange Consortium via the PRIDE (Perez-Riverol et al., 2025) partner repository with the dataset identifier PXD081234.

### Glycolate dehydrogenase assay

Glycolate dehydrogenase activity was measured as described in Lord (1972) using 25-50 μg of lysed chloroplast proteins added to 0.2 mL of reaction mixture (40 mM potassium phosphate, pH 8.0, 80 μM DCIP, and 10 mM sodium glycolate). The assays were performed for 10 min at 30 °C and terminated by adding 8 μL of 12 M HCl and incubated on ice for 10 min. The reaction mixtures were centrifuged for 5 min at 20,000 × g at room temperature and the supernatants were incubated with 42 μL of 0.134 M phenylhydrazine for 10 min at room temperature. Glyoxylate phenylhydrazone was quantified by measuring absorbance at 324 nm in a 96-well microplate using a Synergy H1 microplate reader (BioTek).

### Gas exchange and chlorophyll fluorescence measurements

Gas exchange and chlorophyll fluorescence measurements were performed using a LI-6800 portable photosynthesis system (LI-COR) equipped with a fluorometer chamber on the youngest fully expanded leaves of 6-week-old plants. Prior to measurements, the leaves were stabilized at a reference CO_2_ concentration of 450 µmol mol^-1^ and 1250 µmol m^-2^ s^-1^ light. The CO_2_ assimilation response to *C*_i_ (*A*-*C*_i_) was measured at a series of reference CO_2_ concentrations (450, 300, 200, 100, 50, 25, 10, 450, 450, 450, 450, 600, 750, 900, 1200, 1500, 2000 µmol mol^-1^) at constant light of 1250 µmol m^-2^ s^-1^, leaf temperature of 25 °C, and 50% relative humidity. Subsequently, the response of CO_2_ assimilation to light (*A*-*Q*) was measured at a series of light intensities (1250, 900, 500, 200, 150, 100, 75, 50, 30, 0 µmol m^-2^ s^-1^) at constant reference CO_2_ concentration of 450 µmol mol^-1^, leaf temperature of 25 °C, and 50% relative humidity. *V*_CMAX_ and *J*_MAX_ were estimated from *A*-*C*_i_ curves in R 4.3.1 using PhotoGEA package 1.3.2 (Lochocki et al., 2025) based on Farquhar-von-Caemmerer-Berry model of C_3_ photosynthesis (Farquhar et al., 1980). Γ was estimated as the x-intercept of the fitted *A*-*C*_i_ curve at which net CO_2_ assimilation was zero. From *A*-*Q* curves, *A*_SAT_ was estimated by fitting a non-rectangular hyperbola and Φ_CO2_ was estimated as the initial slope by linear regression over low light range (0-150 µmol m^-2^ s^-1^). ETR was corrected for leaf absorptance measured using a CI-710s SpectraVue leaf spectrometer (CID Bio-Science).

### Mesophyll conductance measurements

The youngest fully expanded leaves of 6-week-old plants was placed in a fluorometer chamber attached to a LI-6800 gas-exchange system (LI-COR) with a leaf temperature of 25 °C, CO_2_ sample of 450 µmol mol^-1^, irradiance of 1250 µmol m^-2^ s^-1^, and an [O_2_] of 21%. The leaf was allowed to acclimate until photosynthesis and stomatal conductance were stable approximately 20-30 min.

The gas-exchange system was coupled to a tunable diode laser (TDL—model TGA 200A; Campbell Scientific Inc.) to measure [^12^CO_2_], [^13^CO_2_] and δ^13^C (Barbour et al., 2007; Bickford et al., 2009; Bowling et al., 2003). The TDL was calibrated using a two-point calibration method where a low CO_2_ calibration tank total [CO_2_] 71.13 ppm (^12^CO_2_ 70.38 ppm and ^13^CO_2_ 0.75 ppm), δ^13^C -39.9‰ and a high CO_2_ calibration tank total [CO_2_] 1472.8 ppm (^12^CO_2_ 1457.16 ppm and ^13^CO_2_ 15.64 ppm), δ^13^C -40.5‰ spanning the range of [CO_2_] for the reference and sample of the gas exchange system. The TDL measurement cycle consisted of the low and high calibration tank followed by the reference and sample gas stream from the gas-exchange system with each measured for 30 s. Once the leaf reached steady state, the gas-exchange system was set to log for approximately 30 minutes. The gas exchange and TDL data were processed and analyzed using PhotoGEA (Lochocki et al., 2025) following the steps described in the “Analyzing Mesophyll Conductance Data” article included with PhotoGEA, which is also available online at the PhotoGEA documentation website: https://eloch216.github.io/PhotoGEA/.

Photosynthetic discrimination (δ^13^C) was calculated according to Evans and Von Caemmerer (2013). The ratio of the ^12^CO_2_ mole fraction in the dry air coming into the gas-exchange cuvette over the difference in ^12^CO_2_ mole fractions of air in and out of the cuvette, ξ was calculated according to Evans et al. (1986). The ternary gas correction factor (*t*) was calculated from Farquhar and Cernusak (2012). The CO_2_ compensation point in the absence of day respiration (Γ*) was calculated assuming a Rubisco specificity of 97.3 M M^-1^ (Walker et al., 2013). Mesophyll conductance to CO_2_ diffusion (*g*_m_) was calculated using equation 13 and 22 from Busch et al. (2020), assuming that mitochondrial respiration is isotopically disconnected from the Calvin-Benson-Bassham cycle. Isotopic fraction due to day respiration (*e*\*) was calculated using equation 19 from Busch et al. (2020) because Δ^growth^ was not available. A week after the TDL measurements, dark respiration was estimated for each individual, these measurements were used for *R*_d_ in the equation to estimate *g*_m_. Outliers were removed within each plant using the 1.5 × interquartile range rule.

### Chlorophyll quantification

Leaf discs 0.5 cm in diameter were sampled from the youngest fully expanded leaves and incubated in 1 mL of 100% (v/v) methanol for 10 min at 55 °C. The chlorophyll extracts were centrifuged for 5 min at 20,000 × g at room temperature and the absorbance of the supernatants was measured at 710, 665, and 650 nm. Chlorophyll *a* and chlorophyll *b* were calculated according to Holden (1976).

### Specific leaf area and aboveground biomass measurements

Specific leaf area was determined using the youngest fully expanded leaves by measuring leaf area using a LI-3100 Area Meter (LI-COR), followed by oven-drying at 60 °C for 6 days and weighing the dried leaves. For aboveground biomass measurements, leaf and stem tissues were harvested from 7-week-old plants, oven-dried at 60 °C for 2 weeks, and weighed.

### Statistical analysis

Statistical analyses were performed in R 4.3.1. For each trait, differences among genotypes were tested using the Kruskal-Wallis test or one-way ANOVA depending on the results of the Shapiro-Wilk and Levene’s tests. When one-way ANOVA was used, post hoc comparisons among genotypes were performed using pairwise *t*-tests with Benjamini-Hochberg correction, while comparisons against WT were performed using Dunnett’s test. When the Kruskal-Wallis test was used, post hoc comparisons were performed using Dunn’s test with Benjamini-Hochberg correction.

## Results

### CrGDH expression in the chloroplast of Chlamydomonas cia5 mutant reduces glycolate accumulation

Plant photorespiration uses glycolate oxidase and catalase to convert glycolate to glyoxylate while producing water as a co-product. We have used CrGDH to catalyze this conversion in part because it may use quinone as an electron acceptor thereby preserving redox energy that is lost when oxygen is the electron acceptor (Meacham-Hensold et al., 2026). CrGDH is natively expressed in the mitochondria of the unicellular microalga *C. reinhardtii* (Beezley et al., 1976). To explore the effect of chloroplast expression of CrGDH prior to plant transformation, we first used *C. reinhardtii* as a rapid and robust system for both nuclear and chloroplast genetic manipulation. In *Chlamydomonas*, knockout of the nuclear-encoded *CIA5* gene, a master regulator of both photorespiration and the carbon concentrating mechanism (CCM), results in excessive production of glycolate (Yun et al., 2021). We exploited this phenotype to evaluate whether chloroplast-localized CrGDH could effectively metabolize glycolate.

First, we generated a *cia5* knockout mutant (*cia5* KO) using CRISPR-Cas9 RNP-mediated knock-in (Figure 1a). In this strain, a hygromycin resistance cassette was inserted into the Cas9 target site within the *CIA5* locus via non-homologous end joining, thereby disrupting CIA5 function. Genomic DNA sequencing of the *CIA5* locus confirmed the gene disruption (Figure 1b). Using this *cia5* mutant background, we next generated chloroplast *psbH* knockout lines by chloroplast transformation to create a recipient line for targeted transgene insertion at the *psbH* locus. PsbH is a small phosphoprotein subunit of photosystem II, and in its absence no stable photosystem II accumulates, resulting in loss of photosynthetic capacity (Summer et al., 1997). We subsequently introduced a CrGDH transgene and reintroduced a functional copy of *psbH* at its native locus, thereby restoring photosynthetic capacity (*cia5* KO with CrGDH OE; Figure 1a). Immunoblot analysis confirmed expression of CrGDH in the resulting transformants (Figure 1c). We then assessed glycolate accumulation in these lines relative to the background strains. As shown in Figure 1d, the *cia5* KO with CrGDH OE strain exhibited a 49% reduction in glycolate accumulation at day 5 compared with the *cia5* KO strain, demonstrating that CrGDH functions effectively in the chloroplast to consume glycolate.

**Figure 1.**
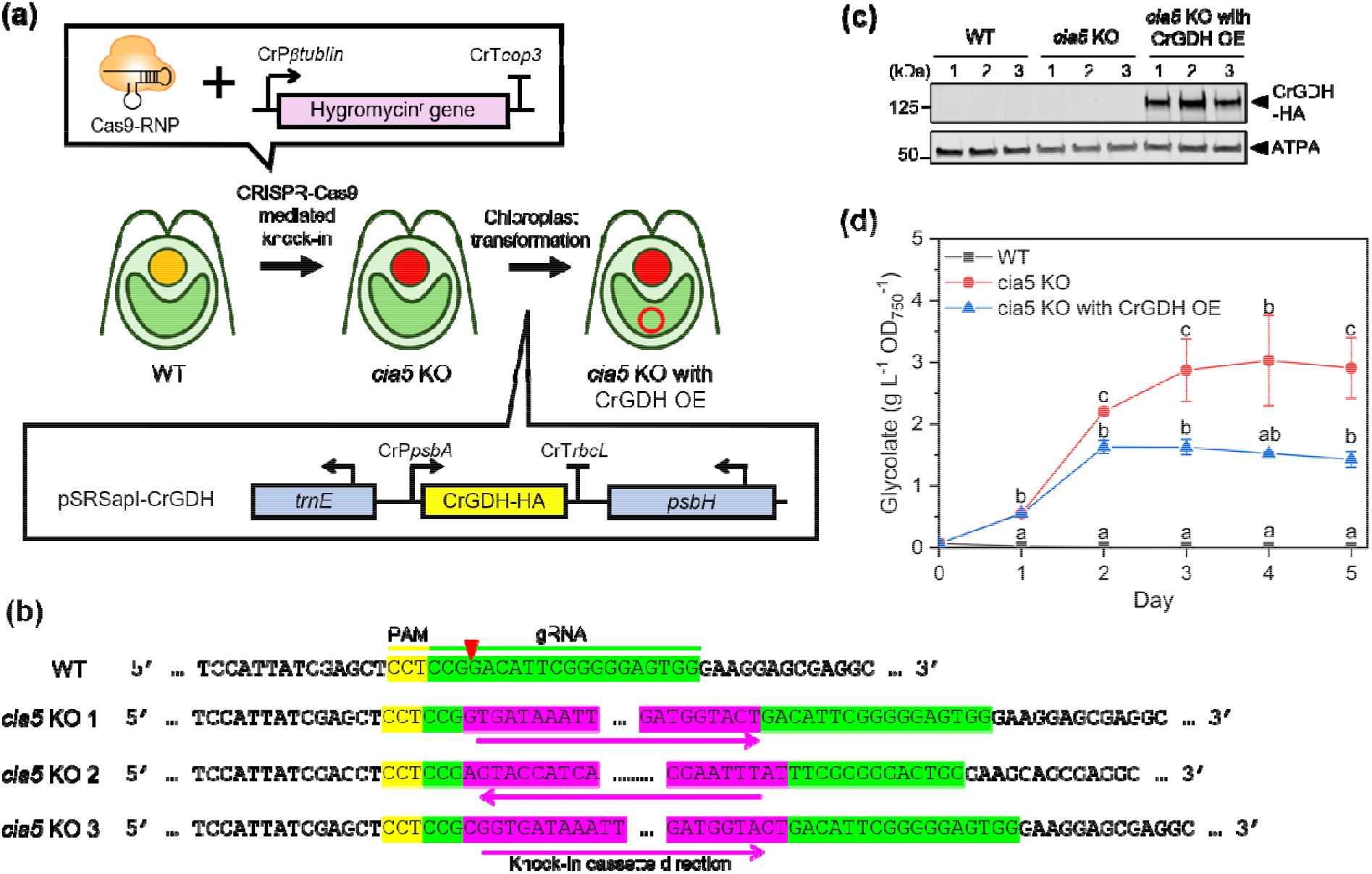
Overexpression of CrGDH in the chloroplast of *Chlamydomonas*. (a) Schematic diagram of generating *cia5* KO and *cia5* KO with CrGDH OE strains. The *cia5* KO strain was generated by CRISPR-Cas9 RNP mediated knock-in using a hygromycin resistance cassette. The *cia5* KO with CrGDH OE strain was subsequently generated by chloroplast transformation with pSRSapI vector carrying *CrGDH-HA* gene under *C. reinhardtii psbA* promoter and *rbcL* terminator. (b) Genomic sequencing confirming disruption of the *CIA5* gene. The wild type (WT) sequence corresponds to CRISPR-Cas9 target region within the second exon of the gene. PAM sequences are highlighted in yellow, gRNA target sequences in green, and the knock-in cassette in magenta. (c) Immunoblot analysis confirming protein expression of CrGDH using anti-HA antibody. Antibody against Alpha subunit of chloroplastic ATP synthase (ATPA) was used as a loading control. (d) Glycolate accumulation in CrGDH overexpressor and parental strains, normalized to cell density (OD_750_). Error bars represent SE and statistical significance was determined by one-way ANOVA followed by pairwise *t*-tests with Benjamini-Hochberg correction or the Kruskal-Wallis test followed by Dunn’s post-doc test with Benjamini-Hochberg correction (n = 3). Different letters indicate significant differences (*P* < 0.05; see supporting data 1 for *P*-values).

### CrGDH was successfully introduced in the tobacco chloroplasts

Based on our findings in *Chlamydomonas*, we expected that expression of CrGDH in tobacco chloroplasts by chloroplast transformation would provide a highly active first step in the photorespiratory bypass by converting glycolate to glyoxylate in the chloroplast. To test this, we assembled the constructs shown in Figure 2a: GFP control vector (pPTS78), CrGDH-GFP fusion vectors for visualizing CrGDH localization (pAPP2881 and pAPP2883), and CrGDH vectors carrying a C-terminal HiBiT tag for assessing GDH function (pAPP2882 and pAPP2884). Constructs pPTS78, pAPP2883, and pAPP2884 were designed for integration of the expression cassettes into the intergenic region between the *trnV* gene and the *3’rps12/7* genes of the Inverted Repeat region of the chloroplast genome and thus are present in two copies per genome. To evaluate any difference in efficacy of CrGDH activity based on copy number, constructs pAPP2881 and pAPP2882 targeted the intergenic region between *rbcL* and *accD* in the Large Single Copy region of the chloroplast genome and are therefore present in only 1 copy per genome. These vectors were introduced into *Nicotiana tabacum* cv. Petit Havana by chloroplast transformation using standard procedures (Bélanger et al., 2023), and homoplasmy was confirmed in the T0 lines after multiple rounds of regeneration by insert sequencing and in the T1 generation by uniform antibiotic resistance in maternally derived seedlings (Supplementary figure S1). The resulting T1 generation was used for all subsequent experiments (Figure 2b).

**Figure 2.**
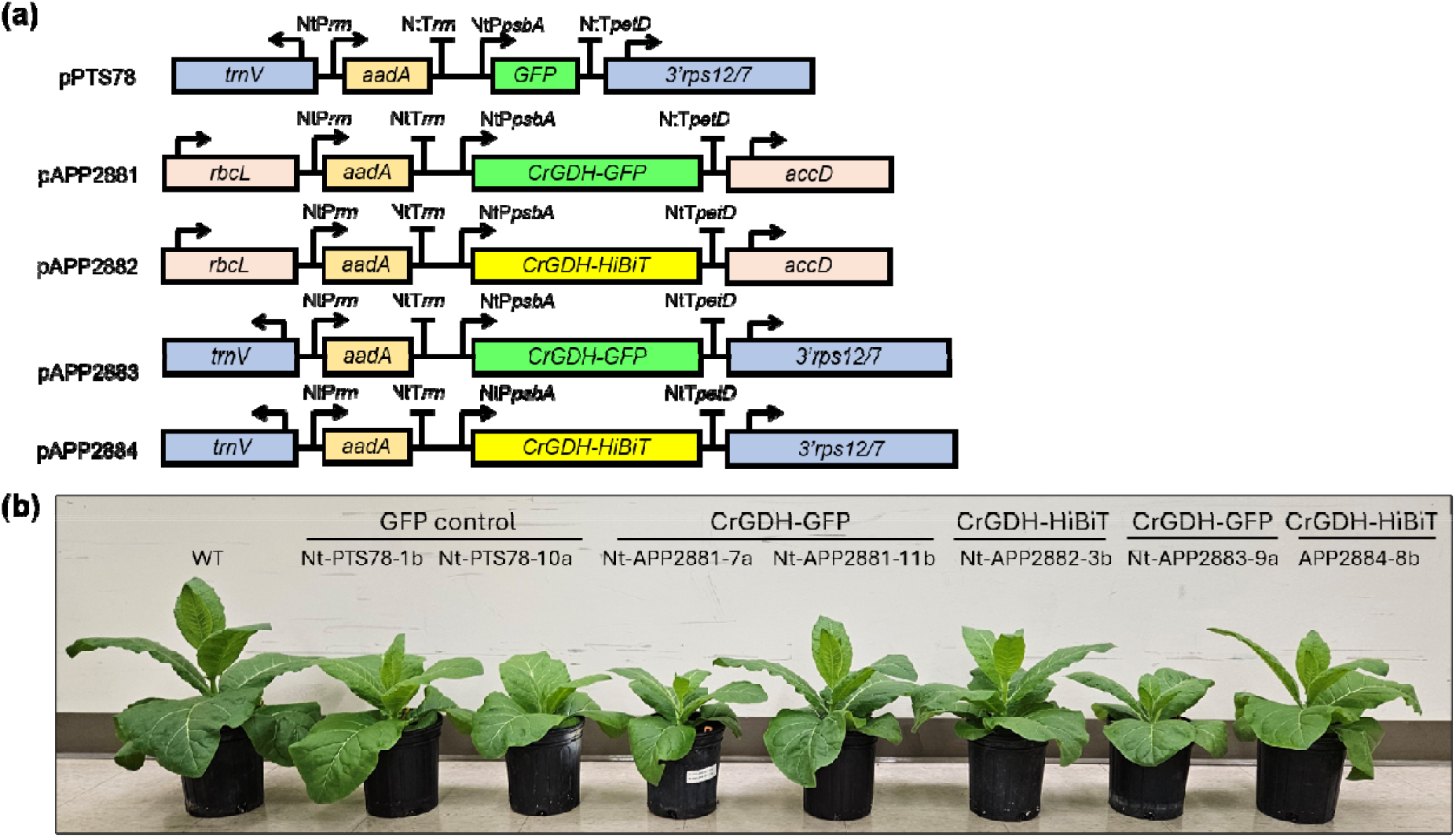
Generation of tobacco chloroplast transformants expressing CrGDH. (a) Vector designs used for chloroplast transformation. Each vector contains the *aadA* gene driven by *N. tabacum* P*rrn* promoter and *E. coli rrnB* terminator, and the gene of interest driven by *N. tabacum psbA* promoter and *petD* terminator. Vectors pPTS78, pAPP2883, and pAPP2884 flank the intergenic region between the *trnV* and *3’rps12/7* genes, whereas pAPP2881 and pAPP2882 flank the intergenic region between *rbcL* and *accD* genes. (b) Representative image of 6-week-old T1 plants generated in this study.

To verify the expression of the intended proteins, we first quantified GFP fluorescence. Quantitative analysis revealed significantly higher fluorescence signals in both GFP control and CrGDH-GFP lines compared to the wild type (WT; Figure 3a), as expected. Confocal microscopy showed that the GFP signal co-localized with chlorophyll fluorescence in both lines, indicating that the proteins were correctly localized within the chloroplasts (Figure 3b).

**Figure 3.**
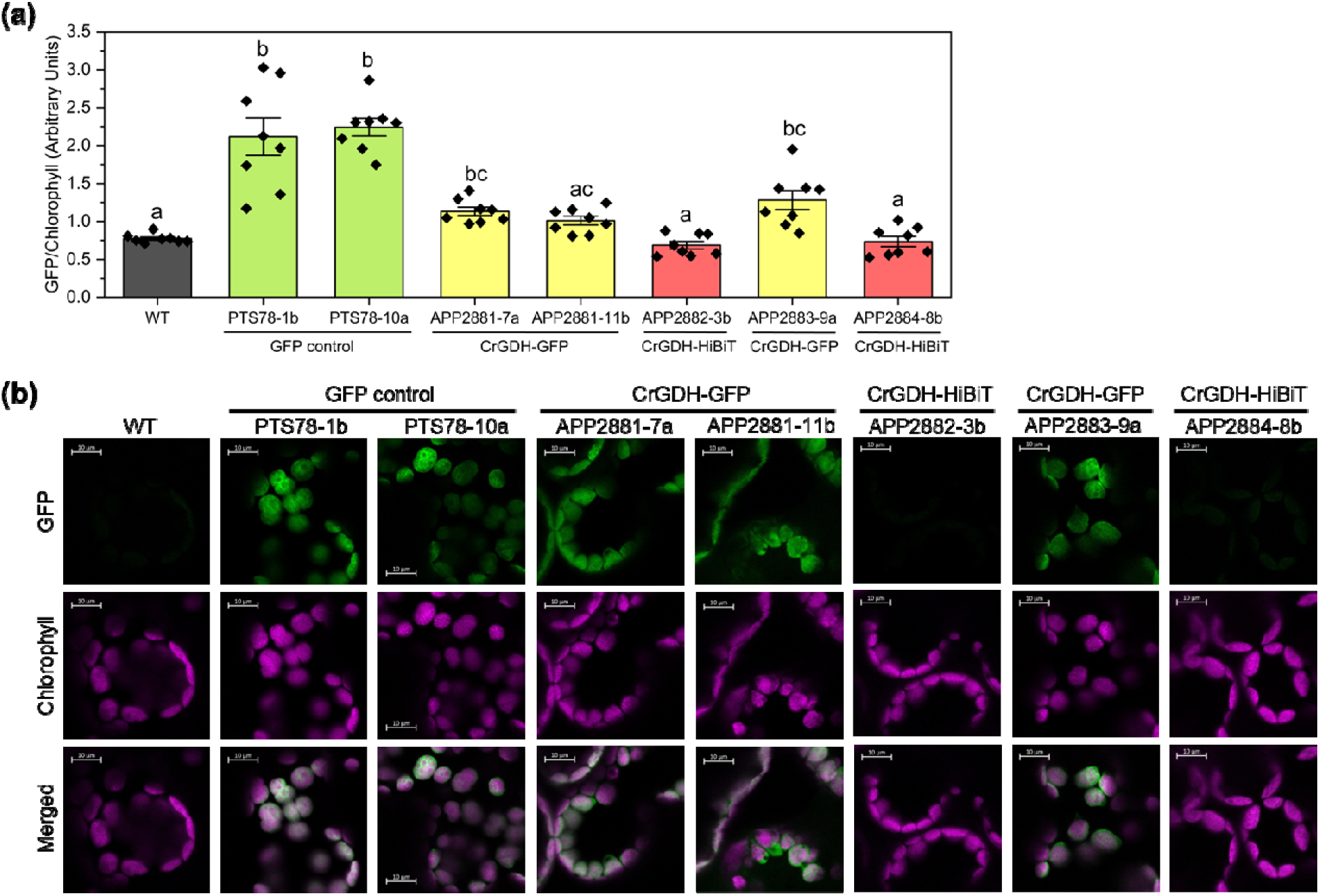
GFP expression in tobacco transgenics. (a) GFP fluorescence normalized to chlorophyll autofluorescence. Data represent means ± standard error. Statistical significance was determined by the Kruskal-Wallis test followed by Dunn’s post-doc test with Benjamini-Hochberg correction (n = 8). Different letters indicate significant differences (*P* < 0.05; see supporting data 1 for *P*-values). (b) Representative confocal microscopy images of GFP fluorescence, chlorophyll autofluorescence, and their merged signal. Scale bar, 10 μm.

We confirmed the accumulation of GFP and CrGDH proteins in the corresponding lines by protein blot analyses using total soluble protein (Figure 4a). To investigate the sub-chloroplastic localization, we isolated chloroplasts and fractionated thylakoids from the selected representative lines. Both GFP-tagged and HiBiT-tagged CrGDH were detected in the thylakoid fractions as well as the chloroplast fractions (Figure 4b).

**Figure 4.**
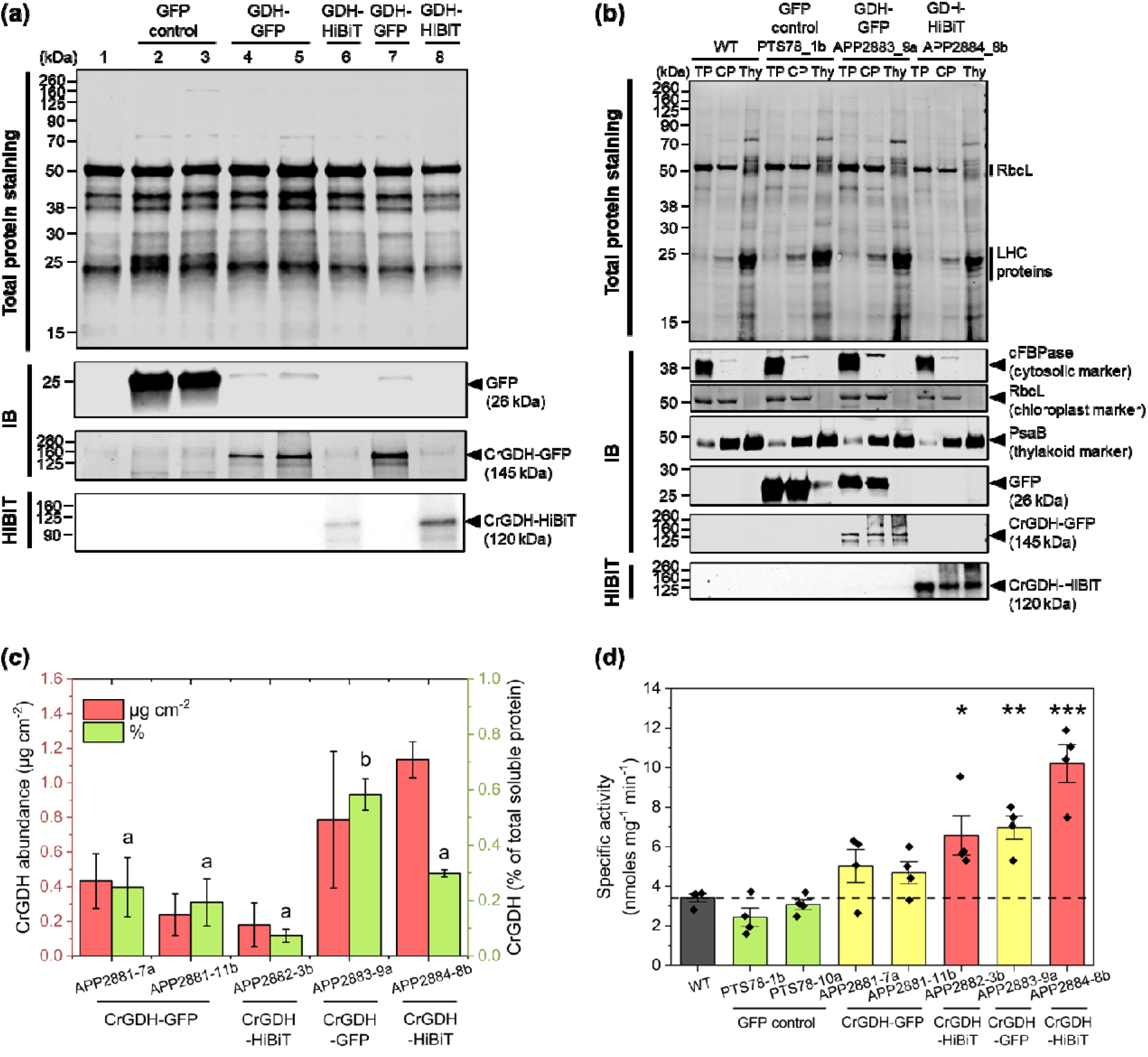
Heterologous CrGDH accumulation and activity in tobacco transgenics. (a) Total protein staining, immunoblotting (IB) with anti-GFP antibody, and HiBiT blotting of total soluble protein extracted from leaves (Lanes: 1, wild type; 2, PTS78-1b; 3, PTS78-10a; 4, APP2881-7a; 5, APP2881-11b; 6, APP2882-3b; 7, APP2883-9a; 8, APP2884-8b). (b) Protein blot analysis of total soluble protein (TP), chloroplast protein (CP), and thylakoid fraction (Thy) isolated from wild type (WT), GFP control (PTS78-1b), GDH-GFP (APP2883-9a), and GDH-HiBiT (APP2884-8b) transgenic plants. Antibodies against Cytosolic fructose-1,6-bisphosphatase (cFBPase), Rubisco large subunit (RbcL), and Photosystem I core PsaB were used as markers for cytosol, chloroplast, and thylakoid, respectively. (c) Absolute CrGDH protein quantification by mass spectroscopy Data represent means ± standard error. Statistical significance was determined by one-way ANOVA followed by pairwise *t*-test with Benjamini-Hochberg correction (n = 3). Different letters indicate significant differences (*P* < 0.05; see supporting data 1 for *P*-values). (d) Specific activity of CrGDH protein in total chloroplast extracts. Data represent means ± standard error. Dashed lines represent the mean value of WT. Statistical analysis was determined by one-way ANOVA followed by Dunnett’s post-hoc test against WT (n = 4). Asterisks indicate significant differences (* *P* < 0.05, ** *P* < 0.01, *** *P* < 0.001; see supporting data 1 for *P*-values).

Protein blot analysis showed the relative abundance of CrGDH among lines carrying either the GFP tag or the HiBiT tag, but it does not provide information on whether the protein accumulates to a physiologically meaningful level across the transgenic lines. To relate the actual amount of accumulated protein to the observed phenotype, we performed mass spectrometry using a spiked-in heavy peptide standard for CrGDH. This absolute quantification further revealed variable accumulation of CrGDH among the transgenic lines, consistent with differences in transgene copy number per chloroplast genome (Figure 4c). CrGDH-HiBiT lines, APP2882-3b (1-copy per chloroplast genome) and APP2884-8b (2-copies per chloroplast genome), accumulated 0.179 and 1.134 μg CrGDH per unit leaf area, corresponding to 0.073% and 0.298% of total soluble protein, respectively. These values correspond to approximately 0.147% and 0.901% of total chloroplast protein based on the estimated chloroplast protein concentration (Supplementary table S5). This accumulation translated into enzymatic activity, as APP2882-3b, APP2883-9a, and APP2884-8b showed significantly higher GDH activity over the background signal seen in WT, with an approximately two to three-fold increase (Figure 4d). The relatively low enzymatic activity observed in APP2881 (1-copy per chloroplast genome) and APP2883 (2-copies per chloroplast genome) lines, despite their detectable protein accumulation, suggests that GFP tag may have a negative effect on enzyme activity, perhaps via steric hindrance of the CrGDH-GFP fusion.

### Overexpression of CrGDH alone is insufficient to improve photosynthesis or biomass in tobacco

Previous studies reported that the transgenic expression of the three subunit EcGDH alone in plants conferred unexpected benefits to photosynthesis and growth in *Arabidopsis*, potato, and *Camelina* (Dalal et al., 2015; Kebeish et al., 2007; Nölke et al., 2014). Such findings were intriguing given that the metabolic fate of glyoxylate produced in the chloroplast by EcGDH remains unclear in the absence of a complete photorespiratory bypass. To determine whether CrGDH induces similar phenotypic changes in tobacco, we evaluated the photosynthetic performance and biomass of the CrGDH single overexpressing lines that were generated via chloroplast transformation.

Photosynthesis was assessed by measuring the light response of CO_2_ assimilation concurrently with chlorophyll fluorescence on the youngest fully expanded leaves of 6-week-old plants under ambient conditions (Figure 5a). In the GFP controls, APP2883-9, and APP2884-8b, operating quantum yield of photosystem II (Φ_PSII_) and electron transport rate (ETR) were reduced compared to WT across all measured light intensities (Figure 5b-c). However, a comparable ETR-to-Net CO_2_ assimilation rate ratio (ETR/*A*_NET_) was observed at most light intensities, except at 500 μmol m^-2^ s^-1^. Apparent quantum yield of CO_2_ assimilation (Φ_CO2_) was also similar between the transgenic lines and WT, indicating that the reduced PSII photochemistry did not substantially alter the coupling between electron transport and carbon fixation (Figure 5d-e). Total chlorophyll content in all transgenic lines, including GFP controls, was approximately 78% of the WT levels (Figure 6a). The chlorophyll *a*/*b* ratio showed no significant difference between the transgenic lines and WT, despite a slight reduction to approximately 90% of WT levels (Figure 6b). These results suggest that the transgenic lines reduced their investment in chlorophyll synthesis to a level that did not compromise the efficiency of CO_2_ assimilation, while the antenna size remained intact.

**Figure 5.**
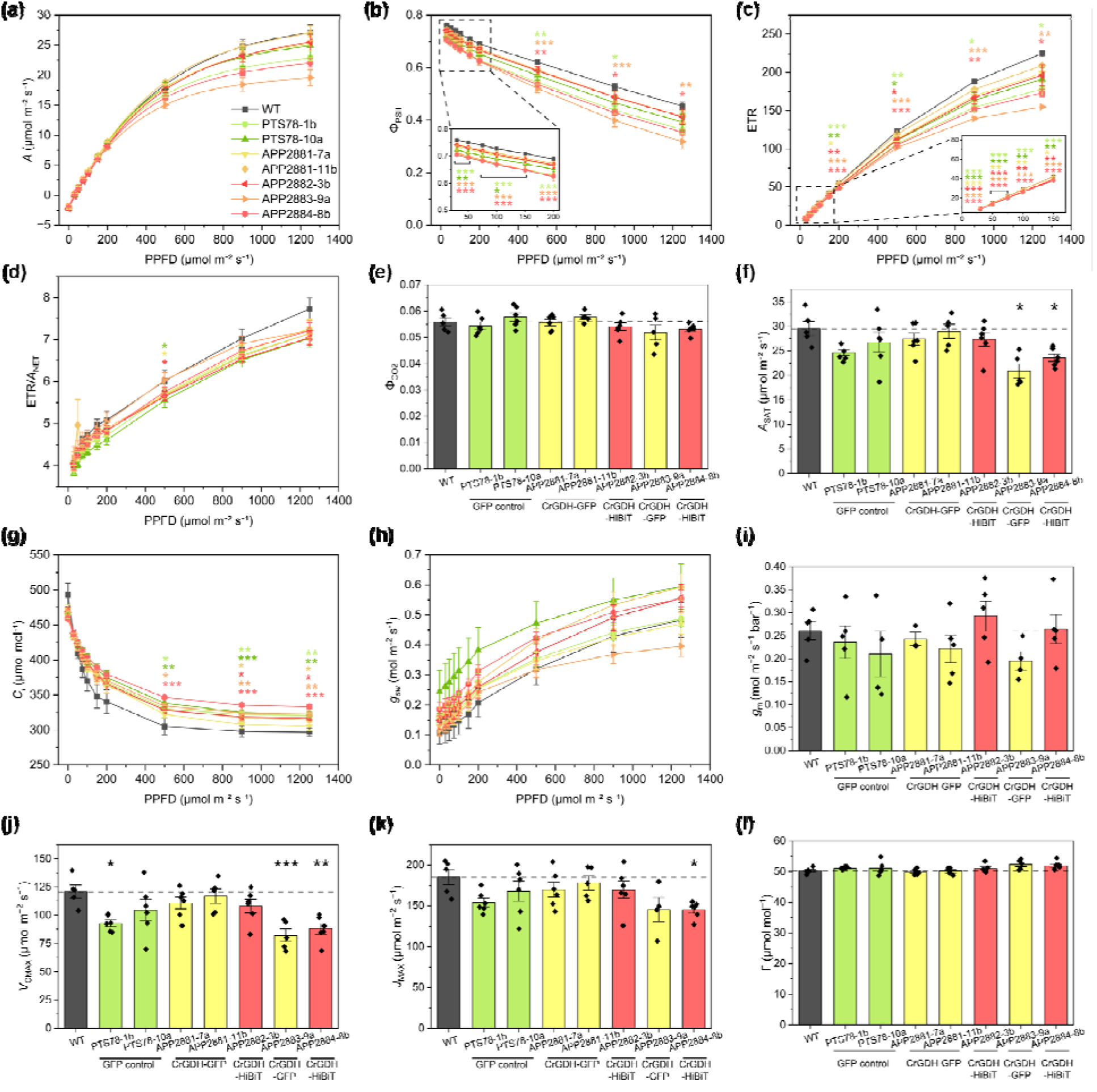
Photosynthetic parameters measured in wild type (WT), GFP controls (PTS78-1b, PTS78-10a), CrGDH-GFP lines (APP2881-7a, APP2881-11b, APP2883-9a), and CrGDH-HiBiT lines (APP2882-3b, APP2884-8b). (a) Light response curves of CO_2_ assimilation (*A*). (b) Operating quantum yield of photosystem II (Φ_PSII_). (c) Electron transport rate (ETR). (d) ETR-to-Net CO_2_ assimilation rate ratio (ETR/*A*_NET_). (e) Apparent quantum yield of photosynthesis (Φ_CO2_). (f) Light saturated CO_2_ assimilation rate at 450 µmol mol⁻¹ CO_2_ (*A*_SAT_). (g) Intercellular CO_2_ concentration (*C*_i_). (h) Stomatal conductance (*g*_sw_). (i) Mesophyll conductance (*g*_m_). (j) Maximum rate of carboxylation (*V*_CMAX_). (k) Maximum rate of electron transport (*J*_MAX_). (l) CO_2_ compensation point (Γ). Data represent means ± standard error. Dashed lines represent the mean value of WT. Statistical analysis was determined by one-way ANOVA followed by Dunnett’s post-hoc test against WT or the Kruskal-Wallis test followed by Dunn’s post-doc test with Benjamini-Hochberg correction (n = 4-6). Asterisks indicate significant differences (* *P* < 0.05, ** *P* < 0.01, *** *P* < 0.001; see supporting data 1 for *P*-values).

**Figure 6.**
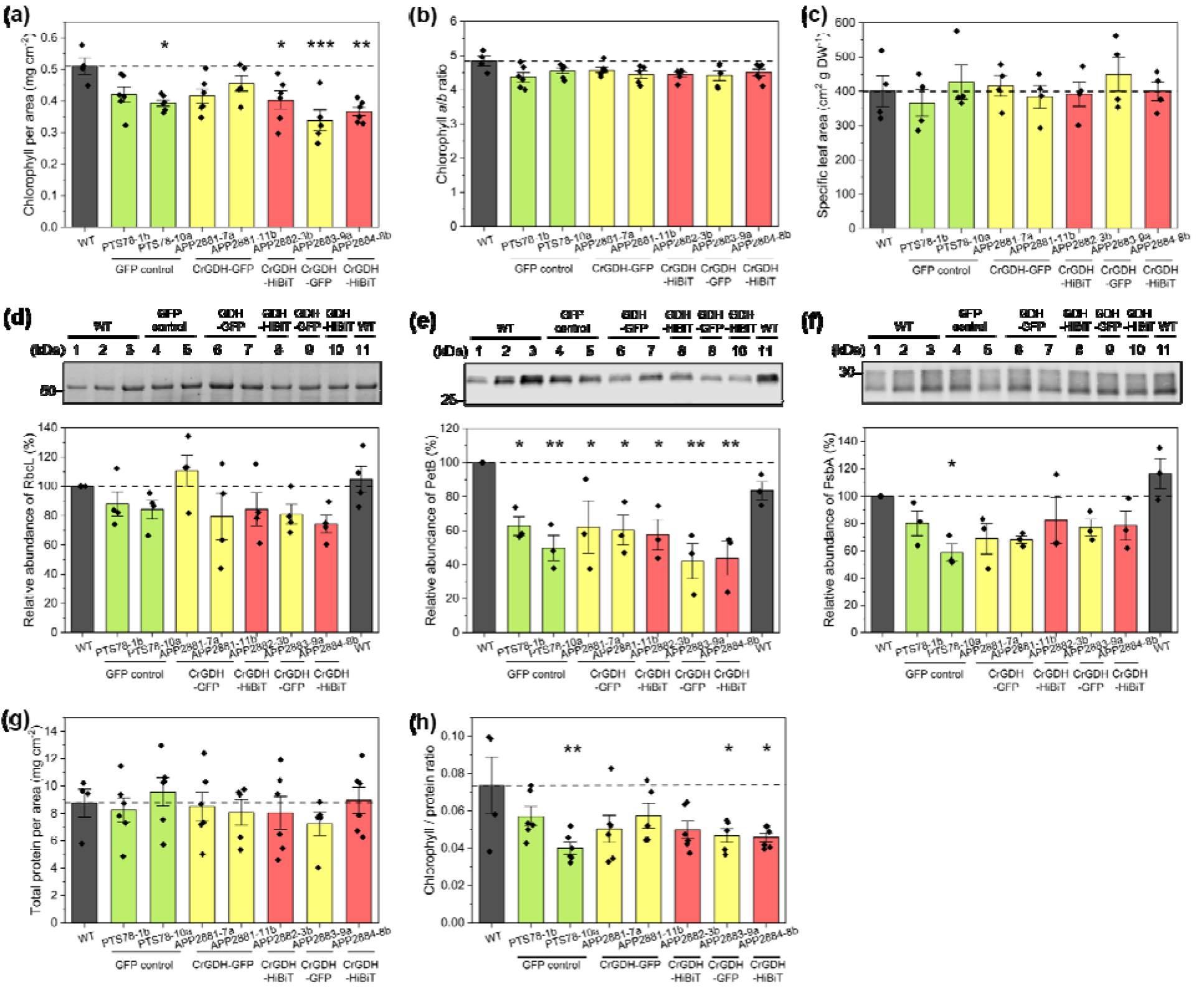
Chlorophyll, specific leaf area, and protein contents in wild type (WT), GFP controls (PTS78-1b, PTS78-10a), CrGDH-GFP lines (APP2881-7a, APP2881-11b, APP2883-9a), and CrGDH-HiBiT lines (APP2882-3b, APP2884-8b). (a) Chlorophyll content per unit area. (b) Chlorophyll *a*/*b* ratio. (c) Specific leaf area. (d) Relative quantification of Rubisco large subunit using total protein staining (RbcL). (e-f) Relative quantification of Cytochrome *b*_6_ (PetB; e) and Photosystem II protein D1 (PsbA; f) using immunoblot analysis. Representative immunoblots probed with PetB and PsbA antibodies, respectively, are shown (Lanes: 1, 25% WT; 2, 50% WT; 3, 100% WT; 4, PTS78-1b; 5, PTS78-10a; 6, APP2881-7a; 7, APP2881-11b; 8, APP2882-3b; 9, APP2883-9a; 10, APP2884-8b; 11, 100% WT). Band intensities were normalized to total protein and expressed as a percentage of 100% WT in lane 3. (g) Total leaf protein content per unit area. (h) Chlorophyll-to-protein ratio. Data represent means ± standard error. Dashed lines represent the mean value of WT. Statistical significance was determined by one-way ANOVA followed by Dunnett’s post-hoc test against WT (n = 3-6). Asterisks indicate significant differences (* *P* < 0.05, ** *P* < 0.01, *** *P* < 0.001; see supporting data 1 for *P*-values).

The light-saturated CO_2_ assimilation rate (*A*_SAT_) was significantly reduced in APP2883-9a and APP2884-8b to 71% and 80% of the WT level, respectively (Figure 5f). Under high-light conditions (500-1250 μmol m^-2^ s^-1^), the transgenic lines maintained higher intercellular CO_2_ concentrations (*C*_i_) with no significant change in stomatal conductance (*g*_sw_), suggesting that the reduction in *A*_SAT_ was not attributed to stomatal limitations (Figure 5g-h). To investigate potential biochemical limitations to the photosynthetic capacity, we estimated maximum rate of carboxylation (*V*_CMAX_) and maximum rate of electron transport (*J*_MAX_) by analyzing CO_2_ assimilation response to *C*_i_ curves corrected for the measured mesophyll conductance (*g*_m_; Figure 5i). *V*_CMAX_ decreased by 77%, 68%, and 73% in PTS78-1b, APP2883-9a, and APP2884-8b, respectively (Figure 5j). Similarly, *J*_MAX_ in APP2884-8b decreased by 78% (Figure 5k). CO_2_ compensation point (Γ) did not differ among the transgenic lines and WT (Figure 5l).

Transgenic lines exhibited significant reductions in Cytochrome *b*_6_ (PetB), a key protein of cytochrome *b*_6_*f* electron transport complex that contributes to *J*_MAX_, with an average decrease of 54% relative to WT (Figure 6e). This is consistent with the lower *J*_MAX_ observed in the transgenic lines, likely reflecting reduced ATP and NADPH production and, consequently, a lower capacity for ribulose-1,5-bisphosphate (RuBP) regeneration. In contrast, abundance of Rubisco large subunit (RbcL) did not differ from WT (Figure 6d). Because *g*_sw_, *g*_m_, and specific leaf area (SLA) were also unchanged in the transgenic lines (Figure 5h, 5i and 6c), limitations in CO_2_ supply or leaf structural traits are unlikely to explain the reduction in the measured *V*_CMAX_. Instead, the lower *V*_CMAX_ may have resulted from a decrease in Rubisco activation state under the reduced energetic status associated with the lower *J*_MAX_. Photosystem II protein D1 (encoded by the *psbA* gene) also showed a decreasing trend, although this reduction was statistically significant only in PTS78-10a (Figure 6f). Overall, the coordinated decreases in light-harvesting and carbon assimilating reactions suggest a coupled reduction in photosynthetic capacity in the transgenic lines. Although total protein content per leaf area was comparable across all lines, chlorophyll-to-protein ratio was generally lower in the transgenic lines (Figure 6g-h). Taken together, these results suggest that heterologous protein accumulation in chloroplasts was associated with adjustments in chlorophyll and photosynthetic protein composition, indicating a partial trade-off between transgene production and native photosynthetic capacity.

The altered photosynthesis impacted aboveground biomass in the transgenic lines (Figure 7). In particular, APP2883-9a, which showed significant reductions in *A*_SAT_, *V*_CMAX_, and *J*_MAX_, displayed a substantial reduction in leaf biomass to 57% of WT. Stem biomass was significantly decreased in PTS78-10a, APP2881-7a, APP2883-9a, and APP2884-8b, with 41, 39, 21, and 45% of WT level, respectively. On average, stem biomass decreased to approximately 55% of WT levels while leaf biomass remained 90% of WT, indicating that the tobacco plants preferentially allocated biomass to leaves over stems to cope with carbon limitations imposed by photosynthetic changes.

**Figure 7.**
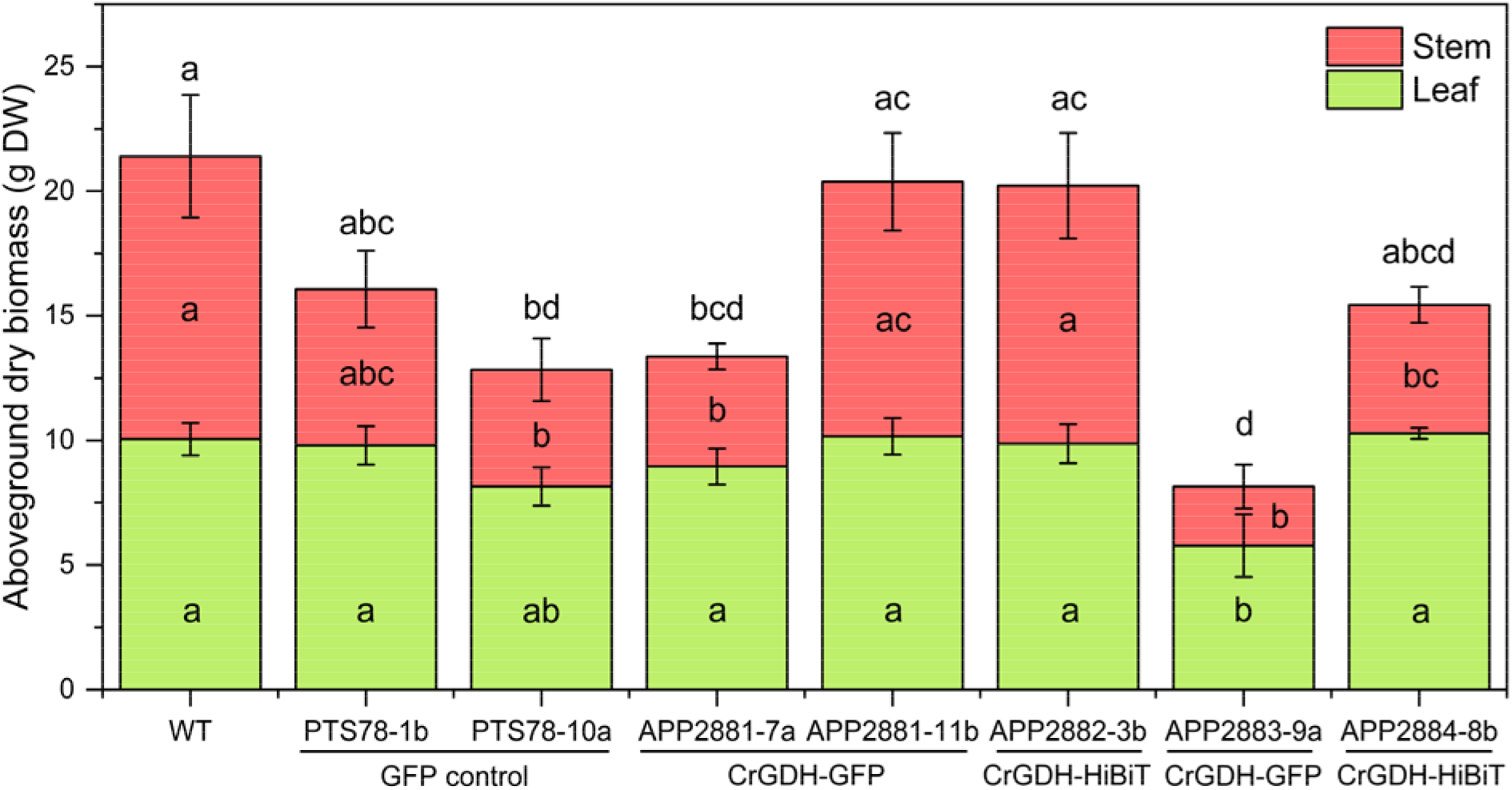
Aboveground dry biomass of wild type (WT), GFP controls (PTS78-1b, PTS78-10a), CrGDH-GFP lines (APP2881-7a, APP2881-11b, APP2883-9a), and CrGDH-HiBiT lines (APP2882-3b, APP2884-8b). Data represent means ± standard error. Statistical significance was determined by one-way ANOVA followed by pairwise *t*-test with Benjamini-Hochberg correction (n = 4-6). Different letters indicate significant differences (*P* < 0.05; see supporting data 1 for *P*-values).

## Discussion

### Chlamydomonas as a rapid platform for testing photorespiratory bypass enzymes

In this study, we introduced CrGDH via chloroplast transformation to establish the first step of photorespiratory bypass in *Chlamydomonas* and tobacco. Although *Chlamydomonas* has a well-developed CCM, a recent study showed that photorespiration still occurs to a substantial extent, particularly under very low CO_2_ concentrations (Dao et al., 2025). Disruption of *CIA5* gene broadly downregulates CCM and photorespiration related genes, leading to excessive glycolate production from Rubisco oxygenation and impaired glycolate metabolism (Dao et al., 2025; Fang et al., 2012). Consequently, glycolate is released into the medium, resulting in carbon loss (Moroney et al., 1986; Yun et al., 2021). In this study, this phenotype was used to assess CrGDH function in the chloroplasts, with reduced glycolate accumulation serving as a clear phenotype readout (Figure 1d). Moreover, the short life cycle of *Chlamydomonas*, together with its accessible genome editing and chloroplast transformation systems (Inckemann et al., 2025; Nievergelt, 2025; Schroda & Remacle, 2022), supports its use as an initial screening platform for photorespiratory bypass enzymes.

### Chloroplast expression of CrGDH in tobacco

In tobacco, the overexpression of CrGDH in transgenic chloroplasts clearly increased the specific activity of the enzyme over the wild-type background (Fig. 4d), indicating success in incorporating this component of the photorespiratory bypass. However, the full potential of chloroplast-expressed CrGDH was difficult to separate from the apparent physiological costs associated with plastid transformation and transgene expression at the time point and growth conditions used in this study. Transgenic lines, including GFP controls, generally showed reduced or WT level photosynthetic performance and aboveground biomass (Figures 5 and 7). In PTS78 lines, this phenotype is consistent with the strong accumulation of GFP, which was readily visible by total protein staining and immunoblotting as an intense band near 25 kDa (Figure 4a). This suggests substantial reallocation of chloroplast resources away from native photosynthetic proteins and chlorophyll (Figure 6). Previous studies have similarly shown that unbalanced accumulation of foreign proteins in chloroplasts can compromise the synthesis of endogenous photosynthetic proteins (Hennig et al., 2007; Oey et al., 2009; Y. Zhang et al., 2023). In contrast, Schmidt et al. (2019) showed no decrease in final biomass of field grown tobacco plants expressing a chloroplast-transgenic cellulase enzyme to greater than 20% total soluble protein. CrGDH accumulation in the APP lines was not excessive despite being expressed under the same regulatory elements as GFP in the PTS78 line (Figure 2 and 4), though biomass and photosynthetic activity were nearly as compromised as in the PTS78 control lines. It is possible that a combination of CrGDH and AadA protein (not tested) in the APP lines together are enough of a physiological burden as the high GFP expression in PTS78 lines is. Y. Zhang et al. (2023) showed that the selectable marker accumulated to up to 7% of total soluble protein, and that the growth phenotype was only observed after removal of the marker. Moreover, lower CrGDH content may not completely reflect the actual biosynthetic cost of its expression, as protein turnover can limit net protein accumulation and increase energetic costs. Unidentified properties of CrGDH coding sequence or protein itself could also have contributed to this effect, potentially through lower translational efficiency, greater structural complexity, or reduced mRNA or protein stability (De Marchis et al., 2012; Hanson et al., 2013). Alternatively, it may be that field-grown tobacco plants compensate over time for recombinant protein expression resulting in biomass accumulation more similar to wild-type plants. It should be noted that chloroplast marker excision technology (Hajdukiewicz et al., 2001; Lutz et al., 2006) could also be used in the future to remove the GFP and aadA selectable marker from chloroplast transgenic plants to address these questions.

These results highlight the importance of balancing transgene expression with chloroplast protein economy and photosynthetic homeostasis, so that the target protein can accumulate to functionally meaningful levels without compromising native chloroplast function. Notably, although APP2882-3a and APP2884-8b harbor identical expression cassettes, their targeted insertion sites into different regions of the chloroplast genome resulted in different transgene copy numbers and consequently, distinct levels of protein accumulation. This suggests that, in addition to the choice of regulatory elements, the strategic selection of genomic integration sites is an important consideration for modulating expression levels. In this regard, APP2882-3a represents a promising candidate for stacking the remaining components of photorespiratory bypass, as it maintains photosynthetic performance comparable to WT while accumulating CrGDH at levels detectable by both protein blot and enzymatic assay (Figure 4-7).

### Estimated chloroplastic abundance of CrGDH

In this study, tobacco transgenic lines accumulated CrGDH at 0.179-1.134 μg cm^-2^ leaf area (Figure 4c). Using reported values of cell number per gram fresh weight, chloroplast number per cell, chloroplast volume, and stromal protein concentration as a proxy for chloroplast protein concentration, CrGDH was estimated to account for 1.47-9.01 μg per mg chloroplast protein (Supplementary table S5). Based on the measured kinetic parameters of CrGDH (Aboelmy & Peterhansel, 2014), these amounts correspond to predicted activities of 1.421-8.709 nmol min^-1^ mg^-1^ chloroplast protein. Given that the assay showed a measurable background signal in the controls, the CrGDH-dependent activities observed in Figure 4d are within a plausible range for the estimated enzyme abundance.

Relative to estimates from the literature, where chloroplast transgenic proteins can represent up to 5-15% of the total soluble protein (Hanson et al., 2013), our estimate should be viewed as a conservative, minimal estimate because it reflects the solubilized protein that was purified as described in the methods; however, CrGDH may preferentially be localized to the thylakoid membrane (Figure 4b) and not completely solubilized by our approaches. Nevertheless, it is substantially higher than in plants carrying genetic modification that originated from nuclear genome and remains within a biologically meaningful range. Absolute protein quantification in *Chlamydomonas* has shown that, excluding Rubisco, Calvin-Benson cycle enzymes are present in the chloroplast over a broad range of approximately 2.1-75.1 μM (Hammel et al., 2020). Thus, even our conservative estimate of CrGDH concentrations of 4.9-30.04 μM (Supplementary table S5) is within the range of native chloroplast enzymes. The physiological relevance of this abundance is further supported by simple kinetic considerations. Glycolate has been reported at approximately 0.02 μmol g FW^-1^ in WT *Arabidopsis* leaves under ambient conditions (Pick et al., 2013). If this entire pool is conservatively assumed to reside within the chloroplast, it would correspond to a chloroplastic glycolate concentration of approximately 1.77 mM. Under this assumption, 4.9 μM CrGDH in APP2882-3a would be predicted to consume glycolate at approximately 0.51 mM min^-1^ based on Michaelis-Menten equation, sufficient to process this pool within about 3.5 min if electron acceptor supply is not limited. Although precise kinetics of PLGG1 and BASS6, which mediate chloroplastic glycolate export, are not well established, available *in vivo* evidence indicates that glycolate export from chloroplast is rapid (Pick et al., 2013; South et al., 2017). ^18^O_2_ labeling experiments showed that glycolate labeling in the WT *Arabidopsis* reached a plateau within 30 s (Pick et al., 2013). This suggests that native glycolate export may compete with CrGDH for stromal glycolate in APP2882-3a. However, recent modeling suggests that alternative photorespiratory pathways may remain beneficial when operating in combination with residual native photorespiratory flux (Smith et al., 2025). Together, these estimates suggest that CrGDH abundance itself is unlikely to be a major limitation in the chloroplast transformants and support the idea that sufficient flux could potentially be redirected through a chloroplast-localized photorespiratory bypass with the addition of downstream bypass enzymes.

### Physiological consequences of CrGDH expression

Previous studies of AP3 bypass in tobacco reported higher Φ_CO2_, raising the possibility that CrGDH may confer a physiological benefit beyond simply diverting glycolate metabolism (Cavanagh et al., 2022; South et al., 2019). The electron acceptor of CrGDH expressed in the chloroplast is unknown. However, based on its reported membrane association and physiological evidence linking mitochondrial CrGDH activity to O_2_ consumption through electron transport chain (Aboelmy & Peterhansel, 2014; Paul et al., 1975; Stabenau & Winkler, 2005), its detection in the thylakoid enriched fraction is consistent with a potential interaction with thylakoid associated electron acceptors. If CrGDH uses plastoquinone, which is abundant in thylakoids, as an electron acceptor, and the resulting electrons contribute to proton accumulation through the Q cycle, the theoretical maximum energetic return would be approximately 0.6 ATP per glycolate (Meacham-Hensold et al., 2026). Such an energetic contribution could improve net CO_2_ assimilation at low irradiance during dusk, dawn, shading or on cloudy days and thereby increase Φ_CO2_ and might not be evident consistently in high light conditions. A statistically significant increase in Φ_CO2_was not consistently observed in AP3 potato (Meacham-Hensold et al., 2024), and AP3 rice has not yet been examined in this context (X. Chen et al., 2026). This suggests that any benefit of CrGDH may be condition dependent and not universally detectable. In this study, overexpression of CrGDH did not increase Φ_CO2_ (Figure 5e) possibly because it comprises a partial bypass that lacks the additional enzymes required to further metabolize glyoxylate, resulting in glyoxylate accumulation in chloroplasts. Glyoxylate has been reported to inhibit Rubisco activation and RUBP regeneration in chloroplasts (Campbell & Ogren, 1990; Chastain & Ogren, 1989; Mulligan et al., 1983), its negative effects may outweigh any energetic advantage derived from electron transfer.

In contrast, overexpression of EcGDH in *Arabidopsis*, potato, and *Camelina* was reported to result in increased photosynthesis and biomass, despite the incompleteness of the bypass (Dalal et al., 2015; Kebeish et al., 2007; Nölke et al., 2014). Feeding ^14^C-glycolate to chloroplast extracts from *Arabidopsis* transgenic lines resulted in significantly greater CO_2_ release than in the WT, indicating that glyoxylate generated by EcGDH can be further converted to CO_2_ via an as-yet unidentified route (Blume et al., 2013; Kebeish et al., 2007). This CO_2_ release may contribute to improved photosynthetic efficiency. However, such benefits were not observed in CrGDH-expressing tobacco examined here, despite both enzymes generating glyoxylate. The potential advantage of CrGDH may have been masked by toxicity associated with glyoxylate and by the reduced photosynthetic capacity observed in the chloroplast transgenic lines (Figure 5-6). In addition, the two GDHs differ in electron acceptor specificity and therefore in their energetic and redox consequences. EcGDH has been proposed to use NAD and/or NADP as electron acceptors in the chloroplast, potentially producing one NAD(P)H per glycolate (Kebeish et al., 2007). In contrast, if CrGDH donates electrons to the plastoquinone pool, the resulting energy may not be fully recovered as ATP, because the proton motive force can also be dissipated through photoprotective processes depending on environmental conditions (Shikanai, 2023). Conversely, NAD(P)H generated by EcGDH may represent a more direct metabolic advantage under conditions in which stromal reducing power is limiting (Fukuda et al., 2023). This advantage would depend on chloroplast redox status, because additional NAD(P)H may be less beneficial, or even detrimental, when the stroma is already highly reduced and oxidized NAD(P) is limiting (Krämer & Kunz, 2021).

The results of the current study indicate that the physiological consequences of CrGDH expression cannot be interpreted solely from its catalytic properties but instead reflect the balance between its potential energetic benefit and multiple physiological constraints of the transgenic lines. A follow-up evaluation of the functional benefit of CrGDH will require evaluation of a complete bypass that includes additional enzymes to process glyoxylate and sustain flux to phosphoglycerate. If CrGDH can support bypass flux in this context, CrGDH could provide an engineering advantage over EcGDH because it is encoded as a single polypeptide, whereas EcGDH requires the introduction and coordinated expression of three separate subunits in the chloroplast. Importantly, if the inhibitory phenotype associated with chloroplast expression of CrGDH can be rescued by introduction of the downstream bypass components, this could serve as a useful phenotypic indicator for identifying transformants in which the complete pathway is functionally established.

## Acknowledgements

We thank J. Bui, Y. Zhu, B. Hajyousif, R. Porwancher, and S. Leon for assistance with plant care and management, tissue harvesting, and data collection. JMS thanks Scout Dour and Kara Boltz for cloning, and Leila Pazouka for tobacco chloroplast insert sequence analysis, respectively. This work was supported by the Realizing Increased Photosynthetic Efficiency (RIPE) project, funded by Gates Agricultural Innovations.

