## Supplementary Figures for "Chloroplast expression of *Chlamydomonas* glycolate dehydrogenase en route to an improved photorespiratory bypass"

**Supplementary figure 1.** Homoplasmy and maternal inheritance of tobacco chloroplast transformants. T1 seeds from self-pollinated T0 plants were germinated on MS media with 500 ug/mL spectinomycin. Chloroplast transformed lines are uniformly green indicating uniform antibiotic resistance and homoplasmy of T1 seedlings, while lack of segregating phenotype confirms maternal inheritance. As control, wild-type tobacco is uniformly bleached on spectinomycin indicating antibiotic sensitivity.

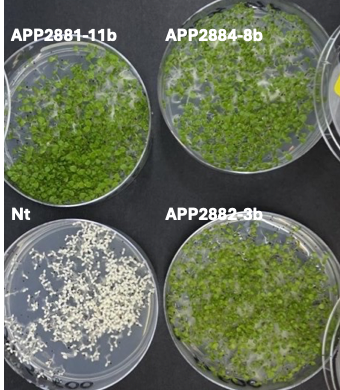

**Supplementary figure 2.** Evaluation of candidate peptides in APP2881-7a and wild type samples by PRM. Eight peptides were assessed for their performance using APP2881-7a as a positive control and wild-type (WT) samples as negative controls. Peptide performance was evaluated based on chromatographic peak area intensity and consistency. Most responsive and specific peptides were subsequently shortlisted for PRM assay development.

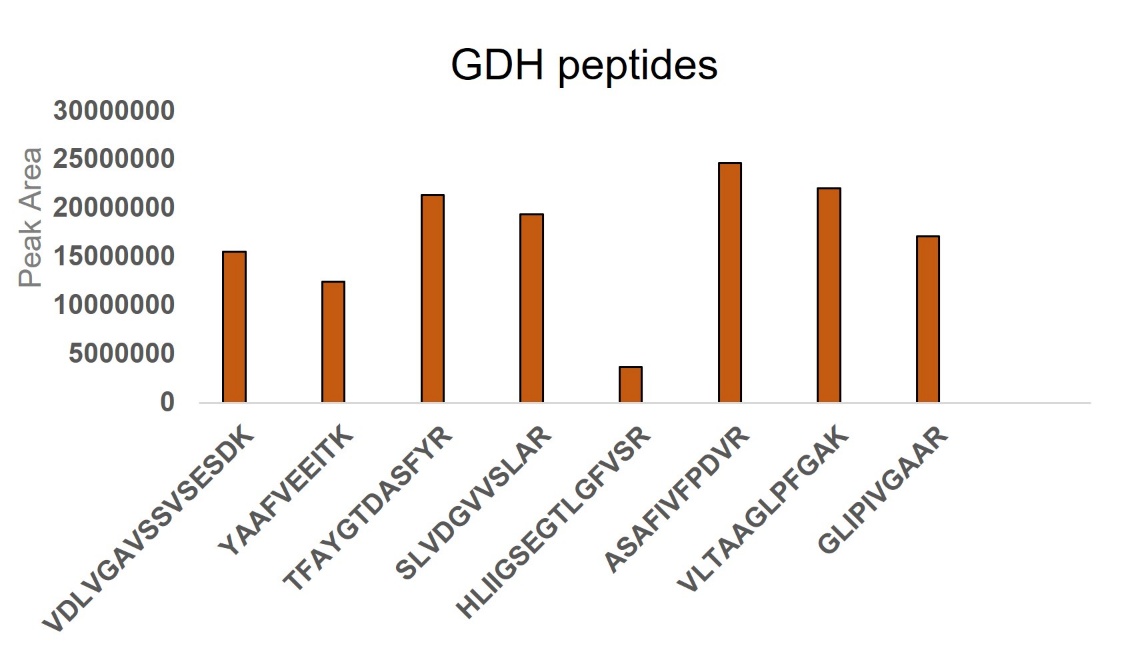
