## Supplementary Tables for "Chloroplast expression of *Chlamydomonas* glycolate dehydrogenase en route to an improved photorespiratory bypass"

**Supplementary table 1.** Primers used in this study

| **No** | **Primer name** | **Sequence (5’-3’)** | **Note** |
| --- | --- | --- | --- |
| 1 | Hyg Insert-Forward | atgattccgctccgtgtaaatg | Amplification of hygromycin resistance cassette for knock-in |
| 2 | Hyg insert-Reverse | agtaccatcaactgacgttacattc |  |
| 3 | CIA5 confirm-Forward | GCACACCCCGCTCTCTTGTT | Colony PCR screening of *cia5* knock-in candidates |
| 4 | CIA5 confirm-Reverse | CCGGCCGGCAAGTGATAGG |  |
| 5 | psbH-down-Forward | TGGGGATGTCAATGCTCCGT | PCR screening of *psbH* knockout candidates |
| 6 | psbH-up-Reverse | ACAAGAAGCAACCCCTTGACGA |  |

**Supplementary table 2.** Skyline generated isolation/transition list for GDH peptides. The table summarizes the candidate peptides, the precursor ion mass-to-charge ratio (m/z), charge states (CS), and polarity.

| **Peptide** | **Mass [m/z]** | **CS [z]** | **Polarity** |
| --- | --- | --- | --- |
| GPASPSSLEQQTR (light) | 679.338955 | 2 | Positive |
| GPASPSSLEQQTR (light) | 453.228395 | 3 | Positive |
| QVAQVAVQQSTQQAVK (light) | 856.968125 | 2 | Positive |
| QVAQVAVQQSTQQAVK (light) | 571.647842 | 3 | Positive |
| VDLVGAVSSVSESDK (light) | 746.380485 | 2 | Positive |
| VDLVGAVSSVSESDK (light) | 497.922749 | 3 | Positive |
| YAAFVEEITK (light) | 585.805696 | 2 | Positive |
| YAAFVEEITK (light) | 390.87289 | 3 | Positive |
| TFAYGTDASFYR (light) | 699.819866 | 2 | Positive |
| TFAYGTDASFYR (light) | 466.882336 | 3 | Positive |
| VHNEDEVR (light) | 499.238512 | 2 | Positive |
| VHNEDEVR (light) | 333.161433 | 3 | Positive |
| LQVPITFR (light) | 487.295102 | 2 | Positive |
| LQVPITFR (light) | 325.19916 | 3 | Positive |
| AAGTSLSGQAITDSVLIK (light) | 866.477991 | 2 | Positive |
| AAGTSLSGQAITDSVLIK (light) | 577.987753 | 3 | Positive |
| NFTVHGDGSVITVEPGLIGGEVNR (light) | 1234.132427 | 2 | Positive |
| NFTVHGDGSVITVEPGLIGGEVNR (light) | 823.09071 | 3 | Positive |
| VVFVDGTVLDTADPNSC[+57.021464]TAFMK (light) | 1194.066518 | 2 | Positive |
| VVFVDGTVLDTADPNSC[+57.021464]TAFMK (light) | 796.380104 | 3 | Positive |
| SLVDGVVSLAR (light) | 558.324588 | 2 | Positive |
| SLVDGVVSLAR (light) | 372.55215 | 3 | Positive |
| C[+57.021464]TTGYSLNALVDFPVDNPIEIIK (light) | 1290.156722 | 2 | Positive |
| C[+57.021464]TTGYSLNALVDFPVDNPIEIIK (light) | 860.44024 | 3 | Positive |
| HLIIGSEGTLGFVSR (light) | 793.438472 | 2 | Positive |
| HLIIGSEGTLGFVSR (light) | 529.29474 | 3 | Positive |
| ATYNTVPEWPNK (light) | 710.348791 | 2 | Positive |
| ATYNTVPEWPNK (light) | 473.901619 | 3 | Positive |
| ASAFIVFPDVR (light) | 611.334955 | 2 | Positive |
| ASAFIVFPDVR (light) | 407.892396 | 3 | Positive |
| AAC[+57.021464]TGASVLR (light) | 503.260933 | 2 | Positive |
| AAC[+57.021464]TGASVLR (light) | 335.843047 | 3 | Positive |
| NETSVDAVELFDR (light) | 747.857177 | 2 | Positive |
| NETSVDAVELFDR (light) | 498.90721 | 3 | Positive |
| EC[+57.021464]ENNEDMMR (light) | 664.239211 | 2 | Positive |
| EC[+57.021464]ENNEDMMR (light) | 443.1619 | 3 | Positive |
| GC[+57.021464]DPMAAALLIEC[+57.021464]R (light) | 788.867653 | 2 | Positive |
| GC[+57.021464]DPMAAALLIEC[+57.021464]R (light) | 526.247528 | 3 | Positive |
| GQDEAALQSR (light) | 537.762351 | 2 | Positive |
| GQDEAALQSR (light) | 358.843993 | 3 | Positive |
| VLTAAGLPFGAK (light) | 572.839873 | 2 | Positive |
| VLTAAGLPFGAK (light) | 382.229008 | 3 | Positive |
| AAQPMAIDAYPFHHDQK (light) | 970.459609 | 2 | Positive |
| AAQPMAIDAYPFHHDQK (light) | 647.308832 | 3 | Positive |
| GLIPIVGAAR (light) | 483.808376 | 2 | Positive |
| GLIPIVGAAR (light) | 322.874676 | 3 | Positive |
| EPGTSMLIEDVAC[+57.021464]PVDK (light) | 930.939527 | 2 | Positive |
| EPGTSMLIEDVAC[+57.021464]PVDK (light) | 620.96211 | 3 | Positive |
| LADMMIDLIDMFQR (light) | 856.414436 | 2 | Positive |
| LADMMIDLIDMFQR (light) | 571.278716 | 3 | Positive |
| FSDMMEEMC[+57.021464]HLVATK (light) | 914.890468 | 2 | Positive |
| FSDMMEEMC[+57.021464]HLVATK (light) | 610.262737 | 3 | Positive |
| NVAPFVEMEWGNK (light) | 760.863751 | 2 | Positive |
| NVAPFVEMEWGNK (light) | 507.578259 | 3 | Positive |
| AYELMWELK (light) | 591.796817 | 2 | Positive |
| AYELMWELK (light) | 394.86697 | 3 | Positive |
| ALFDPSHTLNPGVILNR (light) | 932.507417 | 2 | Positive |
| ALFDPSHTLNPGVILNR (light) | 622.00737 | 3 | Positive |
| FLKPSPAASPIVNR (light) | 748.932825 | 2 | Positive |
| FLKPSPAASPIVNR (light) | 499.624309 | 3 | Positive |
| C[+57.021464]IEC[+57.021464]GFC[+57.021464]ESNC[+57.021464]PSR (light) | 888.333849 | 2 | Positive |
| C[+57.021464]IEC[+57.021464]GFC[+57.021464]ESNC[+57.021464]PSR (light) | 592.558325 | 3 | Positive |
| QLGPGASEEEK (light) | 572.777667 | 2 | Positive |
| QLGPGASEEEK (light) | 382.187537 | 3 | Positive |
| INTGDLIK (light) | 437.255642 | 2 | Positive |
| INTGDLIK (light) | 291.83952 | 3 | Positive |
| TASGMADWLAANFGVINSNVPR (light) | 1146.065501 | 2 | Positive |
| TASGMADWLAANFGVINSNVPR (light) | 764.379426 | 3 | Positive |
| FLNIVNAMHSVVGSAPLSAISR (light) | 1142.117537 | 2 | Positive |
| FLNIVNAMHSVVGSAPLSAISR (light) | 761.74745 | 3 | Positive |
| ALNAATNHFVPVWNPYMPK (light) | 1085.548759 | 2 | Positive |
| ALNAATNHFVPVWNPYMPK (light) | 724.034931 | 3 | Positive |
| VPAPPAPAAAEASGIPR (light) | 786.430647 | 2 | Positive |
| VPAPPAPAAAEASGIPR (light) | 524.622856 | 3 | Positive |
| VVYMPSC[+57.021464]VTR (light) | 606.299201 | 2 | Positive |
| VVYMPSC[+57.021464]VTR (light) | 404.535226 | 3 | Positive |
| MMGPAASDTETAAVHEK (light) | 873.395286 | 2 | Positive |
| MMGPAASDTETAAVHEK (light) | 582.599283 | 3 | Positive |
| AGYEVIIPEGVASQC[+57.021464]C[+57.021464]GMMFNSR (light) | 1288.573662 | 2 | Positive |
| AGYEVIIPEGVASQC[+57.021464]C[+57.021464]GMMFNSR (light) | 859.384866 | 3 | Positive |
| GAELEAALLK (light) | 507.795132 | 2 | Positive |
| GAELEAALLK (light) | 338.865846 | 3 | Positive |
| IPIVIDTSPC[+57.021464]LAQVK (light) | 827.465841 | 2 | Positive |
| IPIVIDTSPC[+57.021464]LAQVK (light) | 551.979652 | 3 | Positive |
| SQISEPSLR (light) | 508.772188 | 2 | Positive |
| SQISEPSLR (light) | 339.517217 | 3 | Positive |
| FALYEPVEFIR (light) | 692.368995 | 2 | Positive |
| FALYEPVEFIR (light) | 461.915089 | 3 | Positive |
| DQVAIHVPC[+57.021464]SSK (light) | 670.834994 | 2 | Positive |
| DQVAIHVPC[+57.021464]SSK (light) | 447.559088 | 3 | Positive |
| MGIEESFAK (light) | 506.244417 | 2 | Positive |
| MGIEESFAK (light) | 337.832037 | 3 | Positive |
| LAGLC[+57.021464]ANEVVPSGIPC[+57.021464]C[+57.021464]GMAGDR (light) | 1202.547447 | 2 | Positive |
| LAGLC[+57.021464]ANEVVPSGIPC[+57.021464]C[+57.021464]GMAGDR (light) | 802.034057 | 3 | Positive |
| FPELTGASLQHLNLPK (light) | 882.985786 | 2 | Positive |
| FPELTGASLQHLNLPK (light) | 588.992949 | 3 | Positive |
| TC[+57.021464]EMSLSNHAGINFR (light) | 868.895788 | 2 | Positive |
| TC[+57.021464]EMSLSNHAGINFR (light) | 579.599617 | 3 | Positive |
| GLVYLVDEATAPK (light) | 688.377017 | 2 | Positive |
| GLVYLVDEATAPK (light) | 459.25377 | 3 | Positive |
| TAGSVSGWR (light) | 460.732866 | 2 | Positive |
| TAGSVSGWR (light) | 307.491002 | 3 | Positive |

**Supplementary table 3.** List of peptide transitions used for Parallel Reaction Monitoring (PRM) assay development. The table summarizes the candidate peptides evaluated during assay optimization to identify the most suitable surrogate peptides for targeted quantification. For each peptide, the precursor ion m/z value, product ion m/z value, charge state and retention time are provided. These transitions were screened based on detectability, and signal intensity and stability prior to selection of the final PRM assay peptides.

| **Peptide** | **Precursor m/z** | **Precursor Charge** | **Product m/z** | **Product Charge** | **Fragment Ion** | **Retention Time** |
| --- | --- | --- | --- | --- | --- | --- |
| VDLVGAVSSVSESDK | 746.380485 | 2 | 1392.68528 | 1 | y14 | 13.28 |
| VDLVGAVSSVSESDK | 746.380485 | 2 | 1392.68528 | 1 | y14 | 13.57 |
| VDLVGAVSSVSESDK | 746.380485 | 2 | 1065.505859 | 1 | y11 | 13.28 |
| VDLVGAVSSVSESDK | 746.380485 | 2 | 1065.505859 | 1 | y11 | 13.57 |
| VDLVGAVSSVSESDK | 746.380485 | 2 | 1008.484395 | 1 | y10 | 13.28 |
| VDLVGAVSSVSESDK | 746.380485 | 2 | 1008.484395 | 1 | y10 | 13.57 |
| VDLVGAVSSVSESDK | 746.380485 | 2 | 937.447282 | 1 | y9 | 13.28 |
| VDLVGAVSSVSESDK | 746.380485 | 2 | 937.447282 | 1 | y9 | 13.57 |
| VDLVGAVSSVSESDK | 746.380485 | 2 | 283.126836 | 2 | y5 | 13.28 |
| VDLVGAVSSVSESDK | 746.380485 | 2 | 283.126836 | 2 | y5 | 13.63 |
| YAAFVEEITK | 585.805696 | 2 | 1007.540788 | 1 | y9 | 13.22 |
| YAAFVEEITK | 585.805696 | 2 | 1007.540788 | 1 | y9 | 13.28 |
| YAAFVEEITK | 585.805696 | 2 | 936.503674 | 1 | y8 | 13.19 |
| YAAFVEEITK | 585.805696 | 2 | 936.503674 | 1 | y8 | 13.28 |
| YAAFVEEITK | 585.805696 | 2 | 865.46656 | 1 | y7 | 13.19 |
| YAAFVEEITK | 585.805696 | 2 | 865.46656 | 1 | y7 | 13.28 |
| YAAFVEEITK | 585.805696 | 2 | 619.329733 | 1 | y5 | 13.19 |
| YAAFVEEITK | 585.805696 | 2 | 619.329733 | 1 | y5 | 13.28 |
| YAAFVEEITK | 585.805696 | 2 | 490.287139 | 1 | y4 | 13.19 |
| YAAFVEEITK | 585.805696 | 2 | 490.287139 | 1 | y4 | 13.28 |
| YAAFVEEITK | 585.805696 | 2 | 361.244546 | 1 | y3 | 13.19 |
| YAAFVEEITK | 585.805696 | 2 | 361.244546 | 1 | y3 | 13.28 |
| YAAFVEEITK | 585.805696 | 2 | 248.160482 | 1 | y2 | 13.22 |
| YAAFVEEITK | 585.805696 | 2 | 248.160482 | 1 | y2 | 13.28 |
| TFAYGTDASFYR | 699.819866 | 2 | 1150.516364 | 1 | y10 | 12.57 |
| TFAYGTDASFYR | 699.819866 | 2 | 1150.516364 | 1 | y10 | 12.6 |
| TFAYGTDASFYR | 699.819866 | 2 | 1079.47925 | 1 | y9 | 12.57 |
| TFAYGTDASFYR | 699.819866 | 2 | 1079.47925 | 1 | y9 | 12.6 |
| TFAYGTDASFYR | 699.819866 | 2 | 916.415922 | 1 | y8 | 12.63 |
| TFAYGTDASFYR | 699.819866 | 2 | 916.415922 | 1 | y8 | 12.6 |
| TFAYGTDASFYR | 699.819866 | 2 | 859.394458 | 1 | y7 | 12.57 |
| TFAYGTDASFYR | 699.819866 | 2 | 859.394458 | 1 | y7 | 12.6 |
| TFAYGTDASFYR | 699.819866 | 2 | 758.34678 | 1 | y6 | 12.57 |
| TFAYGTDASFYR | 699.819866 | 2 | 758.34678 | 1 | y6 | 12.6 |
| TFAYGTDASFYR | 699.819866 | 2 | 643.319837 | 1 | y5 | 12.57 |
| TFAYGTDASFYR | 699.819866 | 2 | 643.319837 | 1 | y5 | 12.6 |
| TFAYGTDASFYR | 699.819866 | 2 | 572.282723 | 1 | y4 | 12.57 |
| TFAYGTDASFYR | 699.819866 | 2 | 572.282723 | 1 | y4 | 12.6 |
| SLVDGVVSLAR | 558.324588 | 2 | 915.525807 | 1 | y9 | 15.55 |
| SLVDGVVSLAR | 558.324588 | 2 | 915.525807 | 1 | y9 | 15.7 |
| SLVDGVVSLAR | 558.324588 | 2 | 701.43045 | 1 | y7 | 15.52 |
| SLVDGVVSLAR | 558.324588 | 2 | 701.43045 | 1 | y7 | 15.7 |
| SLVDGVVSLAR | 558.324588 | 2 | 644.408986 | 1 | y6 | 15.52 |
| SLVDGVVSLAR | 558.324588 | 2 | 644.408986 | 1 | y6 | 15.7 |
| SLVDGVVSLAR | 558.324588 | 2 | 545.340572 | 1 | y5 | 15.52 |
| SLVDGVVSLAR | 558.324588 | 2 | 545.340572 | 1 | y5 | 15.7 |
| SLVDGVVSLAR | 558.324588 | 2 | 446.272158 | 1 | y4 | 15.52 |
| SLVDGVVSLAR | 558.324588 | 2 | 446.272158 | 1 | y4 | 15.7 |
| HLIIGSEGTLGFVSR | 793.438472 | 2 | 1448.810755 | 1 | y14 | 13.72 |
| HLIIGSEGTLGFVSR | 793.438472 | 2 | 1448.810755 | 1 | y14 | 14.69 |
| HLIIGSEGTLGFVSR | 793.438472 | 2 | 1335.726691 | 1 | y13 | 13.72 |
| HLIIGSEGTLGFVSR | 793.438472 | 2 | 1335.726691 | 1 | y13 | 14.69 |
| HLIIGSEGTLGFVSR | 793.438472 | 2 | 1222.642627 | 1 | y12 | 13.72 |
| HLIIGSEGTLGFVSR | 793.438472 | 2 | 1222.642627 | 1 | y12 | 14.69 |
| HLIIGSEGTLGFVSR | 793.438472 | 2 | 1109.558563 | 1 | y11 | 13.72 |
| HLIIGSEGTLGFVSR | 793.438472 | 2 | 1109.558563 | 1 | y11 | 14.69 |
| HLIIGSEGTLGFVSR | 793.438472 | 2 | 836.462478 | 1 | y8 | 13.72 |
| HLIIGSEGTLGFVSR | 793.438472 | 2 | 836.462478 | 1 | y8 | 14.69 |
| HLIIGSEGTLGFVSR | 793.438472 | 2 | 565.309272 | 1 | y5 | 13.72 |
| HLIIGSEGTLGFVSR | 793.438472 | 2 | 565.309272 | 1 | y5 | 14.69 |
| ASAFIVFPDVR | 611.334955 | 2 | 992.556378 | 1 | y8 | 18.08 |
| ASAFIVFPDVR | 611.334955 | 2 | 992.556378 | 1 | y8 | 18.08 |
| ASAFIVFPDVR | 611.334955 | 2 | 845.487965 | 1 | y7 | 18.08 |
| ASAFIVFPDVR | 611.334955 | 2 | 845.487965 | 1 | y7 | 18.08 |
| ASAFIVFPDVR | 611.334955 | 2 | 732.403901 | 1 | y6 | 18.08 |
| ASAFIVFPDVR | 611.334955 | 2 | 732.403901 | 1 | y6 | 18.08 |
| ASAFIVFPDVR | 611.334955 | 2 | 633.335487 | 1 | y5 | 18.08 |
| ASAFIVFPDVR | 611.334955 | 2 | 633.335487 | 1 | y5 | 18.08 |
| ASAFIVFPDVR | 611.334955 | 2 | 486.267073 | 1 | y4 | 18.08 |
| ASAFIVFPDVR | 611.334955 | 2 | 486.267073 | 1 | y4 | 18.08 |
| ASAFIVFPDVR | 611.334955 | 2 | 274.187366 | 1 | y2 | 18.11 |
| ASAFIVFPDVR | 611.334955 | 2 | 274.187366 | 1 | y2 | 18.08 |
| VLTAAGLPFGAK | 572.839873 | 2 | 932.519993 | 1 | y10 | 12.27 |
| VLTAAGLPFGAK | 572.839873 | 2 | 932.519993 | 1 | y10 | 14.46 |
| VLTAAGLPFGAK | 572.839873 | 2 | 831.472314 | 1 | y9 | 12.27 |
| VLTAAGLPFGAK | 572.839873 | 2 | 831.472314 | 1 | y9 | 14.46 |
| VLTAAGLPFGAK | 572.839873 | 2 | 760.435201 | 1 | y8 | 12.45 |
| VLTAAGLPFGAK | 572.839873 | 2 | 760.435201 | 1 | y8 | 14.46 |
| VLTAAGLPFGAK | 572.839873 | 2 | 689.398087 | 1 | y7 | 12.27 |
| VLTAAGLPFGAK | 572.839873 | 2 | 689.398087 | 1 | y7 | 14.46 |
| VLTAAGLPFGAK | 572.839873 | 2 | 519.292559 | 1 | y5 | 12.27 |
| VLTAAGLPFGAK | 572.839873 | 2 | 519.292559 | 1 | y5 | 14.46 |
| GLIPIVGAAR | 483.808376 | 2 | 796.503949 | 1 | y8 | 9.21 |
| GLIPIVGAAR | 483.808376 | 2 | 796.503949 | 1 | y8 | 14.78 |
| GLIPIVGAAR | 483.808376 | 2 | 683.419885 | 1 | y7 | 9.68 |
| GLIPIVGAAR | 483.808376 | 2 | 683.419885 | 1 | y7 | 14.81 |
| GLIPIVGAAR | 483.808376 | 2 | 374.214643 | 1 | y4 | 9.23 |
| GLIPIVGAAR | 483.808376 | 2 | 374.214643 | 1 | y4 | 14.81 |
| GLIPIVGAAR | 483.808376 | 2 | 398.755612 | 2 | y8 | 9.21 |
| GLIPIVGAAR | 483.808376 | 2 | 398.755612 | 2 | y8 | 14.81 |
| GLIPIVGAAR | 483.808376 | 2 | 342.21358 | 2 | y7 | 9.62 |
| GLIPIVGAAR | 483.808376 | 2 | 342.21358 | 2 | y7 | 14.81 |

**Supplementary table 4.** Heavy and light forms of the surrogate peptide used for absolute quantification. The table lists the details of both the endogenous (light) and isotopically labeled (heavy) versions of the selected surrogate peptide used for PRM-based absolute quantification. The heavy peptide standard was spiked into samples as an internal reference for normalization and determination of absolute abundance.

| **Peptide** | **Precursor Mz** | **Label** | **Precursor Charge** | **Product Mz** | **Product Charge** | **Fragment Ion** |
| --- | --- | --- | --- | --- | --- | --- |
| GLIPIVGAAR | 483.808376 | Light | 2 | 683.419885 | 1 | y7 |
| GLIPIVGAAR | 483.808376 | Light | 2 | 473.283057 | 1 | y5 |
| GLIPIVGAAR | 483.808376 | Light | 2 | 374.214643 | 1 | y4 |
| GLIPIVGAAR | 483.808376 | Light | 2 | 398.755612 | 2 | y8 |
| GLIPIVGAAR | 488.812511 | Heavy | 2 | 693.428154 | 1 | y7 |
| GLIPIVGAAR | 488.812511 | Heavy | 2 | 483.291326 | 1 | y5 |
| GLIPIVGAAR | 488.812511 | Heavy | 2 | 384.222912 | 1 | y4 |
| GLIPIVGAAR | 488.812511 | Heavy | 2 | 403.759747 | 2 | y8 |

**Supplementary table 5**. Estimation of CrGDH abundance in chloroplasts using reported values and values experimentally determined in this study. Row numbers indicate the input parameters used for each calculation. Stromal protein concentration (parameter 10) was used as proxy for total chloroplast protein concentration. g FW, gram fresh weight; g DW, gram dry weight.

| **No** | **Parameter** | **Value** | **Unit** | **Source / Calculation** |
| --- | --- | --- | --- | --- |
| 1 | Cell number per g FW | 4.5×10^6^ | cells g FW^-1^ | Binding (1975) |
| 2 | Chloroplast number per cell | 100 | chloroplasts cell^-1^ | Staub and Maliga (1992) |
| 3 | Chloroplast volume | 25 | μm^3^ chloroplast^-1^ | Newell and Gray (2010) |
|  |  | 2.5×10^-11^ | mL chloroplast^-1^ |  |
| 4 | Chloroplast volume per g fresh weight | 11.25×10^-3^ | mL g FW^-1^ | Calculated using parameters 1, 2, 3 |
| 5 | CrGDH-HiBit protein per g fresh weight (APP2882-3b) | 6.618 | μg g FW^-1^ | In this study |
| 6 | CrGDH-HiBit protein per g fresh weight (APP2884-8b) | 40.559 | μg g FW^-1^ | In this study |
| 7 | CrGDH-HiBit | 120 | kDa | In this study |
| 8 | Estimated CrGDH abundance in the chloroplast (APP2882-3b) | 0.588 | μg μL^-1^ | Calculated using parameters 4, 5, 7 |
|  |  | 4.9 | μM |  |
| 9 | Estimated CrGDH abundance in the chloroplast (APP2884-8b) | 3.605 | μg μL^-1^ | Calculated using parameters 4, 6, 7 |
|  |  | 30.04 | μM |  |
| 10 | Stroma protein concentration | 400 | mg mL^-1^ | Yabuta et al. (2008) |
| 11 | Estimated CrGDH abundance relative to chloroplast protein (APP2882-3b) | 1.47 | μg mg chloroplast^-1^ | Calculated using parameters 8, 10 |
|  |  | 0.147 | % |  |
| 12 | Estimated CrGDH abundance relative to chloroplast protein (APP2884-8b) | 9.013 | μg mg chloroplast^-1^ | Calculated using parameters 9, 10 |
|  |  | 0.901 | % |  |
